# Asymmetry in synaptic connectivity balances redundancy and reachability in the *C. elegans* connectome

**DOI:** 10.1101/2024.03.06.583122

**Authors:** Varun Sanjay Birari, Ithai Rabinowitch

## Abstract

The brain is overall bilaterally symmetrical, but exhibits also considerable asymmetry. While symmetry may endow neural networks with robustness and resilience, asymmetry may enable parallel information processing and functional specialization. How is this functional tradeoff between symmetrical and asymmetrical brain architecture balanced? To address this, we focused on the *C. elegans* connectome, which comprises 99 classes of bilaterally symmetrical neuron pairs. We found symmetry in the number of synaptic partners between neuron class members, but pronounced asymmetry in the identity of these synapses. We developed graph theoretical metrics for evaluating Redundancy, the selective reinforcement of specific neural paths by multiple alternative synaptic connections, and Reachability, the extent of synaptic connectivity of each neuron class. We found Redundancy and Reachability to be stochastically tunable by the level of network asymmetry, driving the *C. elegans* connectome to favor Redundancy over Reachability. These results elucidate fundamental relations between lateralized neural connectivity and function.

## Introduction

Different patterns of synaptic connectivity enable specific modes of neural circuit function. For example, convergent connections may promote the integration of neural information^1,2^, and hierarchical connectivity may afford multi-layered processing^3^. Perhaps the most prominent gross feature of brain structure is its overall bilateral symmetrical organization^4^, evident in its division, in many species, into left and right hemispheres, and the bilateral duplication of entire brain regions, neuron types and synaptic connections. What are the functional implications of bilateral symmetry in the nervous system?

In many animals, symmetrical brain structure corresponds to bilateral symmetry of the body (*e*.*g*., left-right limbs, eyes, lungs), enabling walking and running, the manipulation of large objects, stereoscopic vision and spatial sound localization. Brain symmetry also provides full sensorimotor coverage from both sides of the body, for better detection of threats^5^ and opportunities^6^. At the neural level, symmetrical duplication of brain regions into left-right equivalents may contribute to robust signaling, reducing noise and uncertainty, overcoming errors or functional deficits, and additively increasing output strength.

At the same time, asymmetry also prevails^7,8^, such as in organ position (*e*.*g*., heart, liver), behavior (*e*.*g*., mothers cradling babies predominantly on their left side^9^), brain connectivity^10^ (*e*.*g*., greater regional interconnectivity in the right vs. left hemisphere^11^), and brain function (*e*.*g*., speech vs. prosody^12^). Such lateralization could enhance specialization, by maximizing the use of brain resources and allocating separate functions to distinct parallel left-right regions^13^. A lateralized brain reduces the need for left-right coordination, which saves energy, shortens reaction times and eliminates potential cross-interferences between left and right circuits. Lateralization may also facilitate multi-tasking through parallel left-right processing^14^. Bilateral compartmentalization can thus enable more efficient and less interruptible brain operation, and has been suggested to enhance cognitive abilities^5,15,16^. Conversely, diminished asymmetry has been associated with several disease conditions^17^. The tradeoff between robustness, additivity and increased coverage, afforded by symmetrical neural organization, on the one hand, and specialization, compartmentalization and parallel processing, enabled by brain lateralization, on the other hand, epitomizes a fundamental interplay between structure and function in the brain.

In order to better understand the functional impact of symmetrical vs. asymmetrical neuronal organization on brain operation, we analyzed the connectome, the map of all synaptic connections, of the nematode worm *C. elegans*, focusing on lateralization in chemical synaptic connectivity. The adult hermaphrodite *C. elegans* nervous system contains 198 (out of 302) bilaterally symmetrical neurons^18^, belonging to distinct neuron classes whose left and right members are clearly labeled^19^. We framed the question of network symmetry vs. asymmetry in terms of Redundancy, the selective reinforcement of specific synaptic paths between distinct neuron classes through parallel alternative connections, and Reachability, the extent of connectivity of each neuron class to other neuron classes. We developed novel metrics based on graph theory for comparing between Redundancy and Reachability, making it possible to resolve the functional balance between network symmetry and asymmetry in several connectome networks and in simulated networks.

## Results

### The number of synaptic contacts of each neuron is consistent across connectome networks

In order to examine asymmetrical features of synaptic connectivity in *C. elegans* we constructed connectome networks from 3 available connectome datasets of adult *C. elegans* (see Methods). We labeled these networks CeH0 (a corrected and updated version of the first published *C. elegans* connectome^20^), and CeH2 and CeH3 (more recently mapped connectomes^21^). CeH2 and CeH3 networks do not include neurons from the body and tail, and thus contain fewer neurons (*N* = 180) than CeH0 (*N* = 280). We thus generated, for the purpose of analysis, a subnetwork of CeH0 containing only the neurons included in the CeH2 and CeH3 networks, and named it CeH1. In addition, we restricted most of our analysis to chemical synapses (networks CeH1c, CeH2c and CeH3c), for which more details exist compared to electrical synapses, and we considered only the presence or absence of connections, ignoring their weights (*i*.*e*., the number of synaptic contacts). As in many other nervous systems^22^, the number of synaptic connections of each neuron in *C. elegans* varies according to neuron type. This number is known as the neuron’s degree. Comparing the degree, *d*_*i*_, of each neuron, *i*, across the 3 connectome networks revealed similar degree distributions (Fig. 1A), and strong correlations between the networks (Fig. 1B), suggesting that the degree of a neuron is to a large extent, a fixed feature of neuronal connectivity.

**Figure 1.**
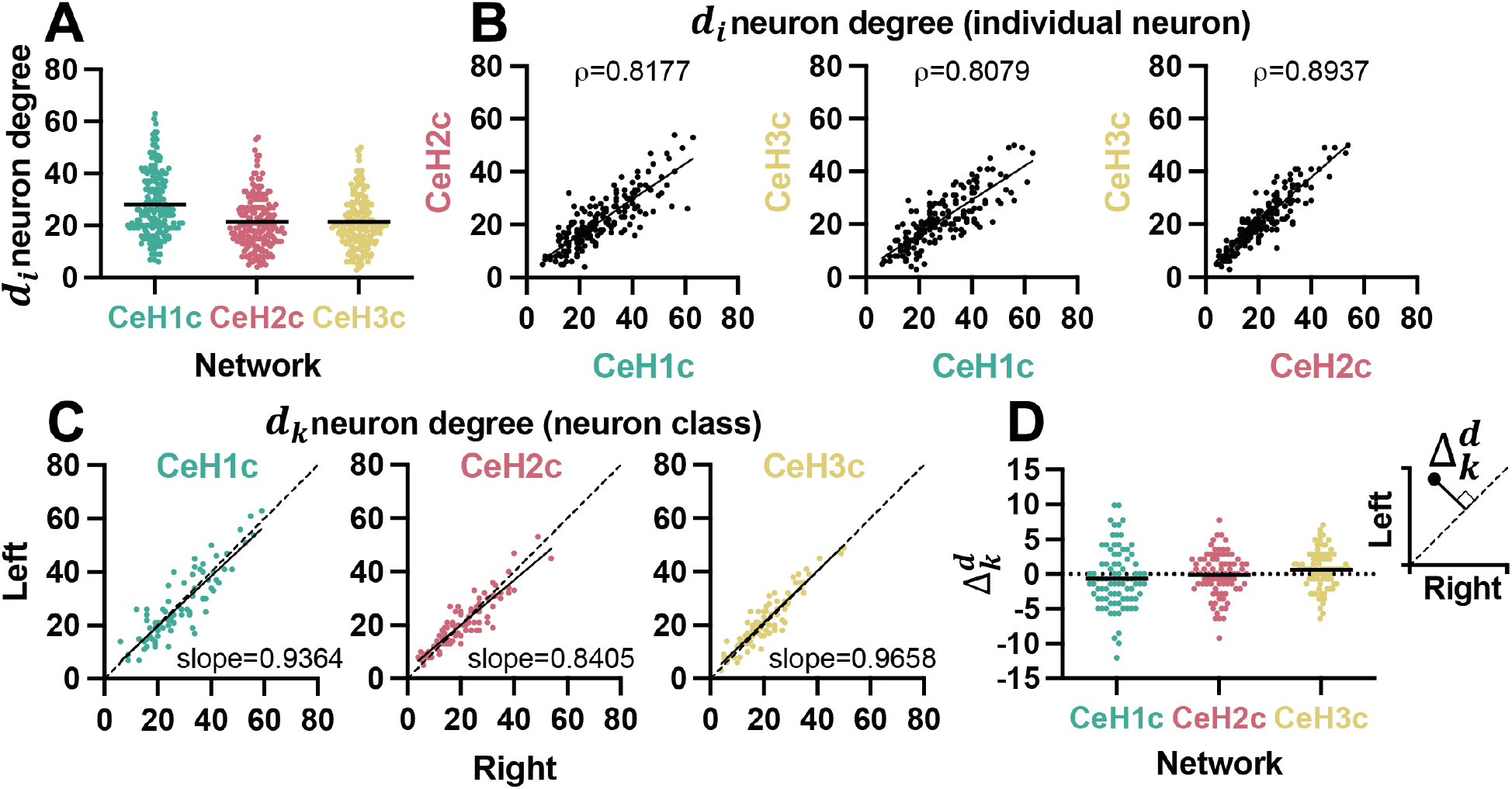
Left-right symmetry in neuron chemical synapse degree. (A) Individual neuron degree, *d*_*i*_, within each connectome network. Dark bars indicate average values. (B) Comparison of *d*_*i*_ between connectome networks. *ρ* - Spearman rank correlation. n=180, p<0.0001 for all plots. (C) Comparisons between left, 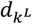, and right, 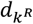, neuron class members for each connectome network. Slopes fitted from simple linear regression. n=83, p<0.0001 for all plots. Dotted diagonal is the identity line. (D) 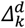, the Euclidean distance of each 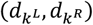coordinate from the symmetry identity line (in C). Inset shows an example neuron class (black dot). Dark bars indicated average values.

### Bilateral neuron pairs exhibit symmetry in the number of synaptic contacts

Given the typical degree of each individual neuron (Fig. 1A), we asked whether the left and right neurons within each neuron class may show similarity in degree. We examined for each neuron class, *k*, the relations between the degree of the left and right members of that class, *k*^*L*^ and *k*^*R*^. We found a strong correlation between 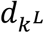and 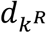(Fig. 1C), with most values lying close to the identity line, representing perfect symmetry (Fig. 1C). This result suggests that in addition to the preserved degree of each individual neuron (Fig. 1B), bilateral neurons within neuron classes tend to maintain degree symmetry. The similar degree of each neuron class member may represent a property of that neuron class.

To evaluate the modest differences in degree that nevertheless exist between left and right class members, we calculated 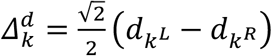, the Euclidean distance of each 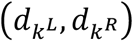 coordinate from the symmetry identity line, as a measure of deviation from class degree symmetry (Fig. 1D, inset). 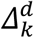showed a comparable distribution between networks, centered close to zero (Fig. 1D), implying no particular overall left or right bias. In addition, 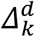values showed no correlation between networks (Fig. S1A), suggesting that the deviations from perfect class symmetry might not be fixed. Finally, 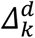seemed to not be strongly dependent on the total degree of the neuron pair, *d*_*k*_ (Fig. S1B), further supporting variable rather than systematic deviations between 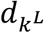and 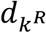. Together, these findings suggest that neuron degree is a consistent and innate property of each neuron class shared by its members, reflecting a fundamental quantitative aspect of network symmetry, including a small stochastic deviation.

### Bilateral neuron pairs show asymmetry in synapse composition

The left-right resemblance in the number of synaptic connections within each neuron class raises the question of whether the composition of these synapses is symmetrical as well. To address this, we examined the synaptic partners of the left versus right members of each neuron class. We considered synapses linking between two contralateral neurons from one class and two contralateral neurons from another class as *paired* synapses. All other synapses that cannot be matched in this way are *unpaired* synapses. For example, in the small model network (Fig. 2A), neuron DL (the left member of neuron class D) participates in 2 synapses, DL-CL and DL-GL. Its contralateral class member neuron, DR (the right member of neuron class D) partakes in 3 synapses, DR-AR, DR-CR and DR-FR. Since CL and CR are contralateral neurons of the same neuron class, C, the DL-CL synapse can be said to be paired with the DR-CR synapse (Fig. 2A, dashed connectors). All other 3 synapses of the D neuron class are unpaired (Fig. 2A, continuous connectors). The fraction of unpaired synapses of this neuron class is 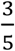(3 unpaired synapses out of a total of 5 synapses for the entire class).

**Figure 2.**
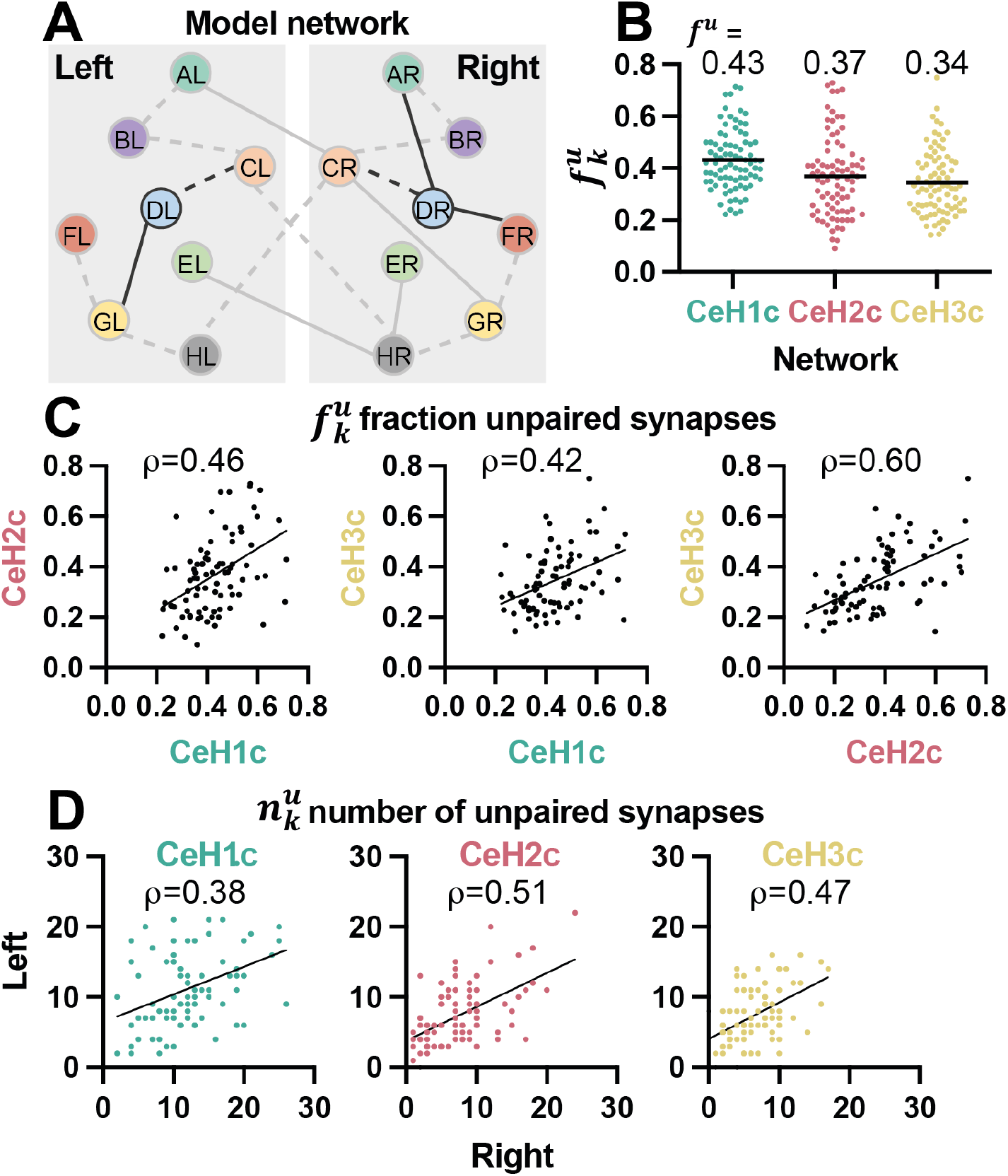
Asymmetry in chemical synaptic composition. (A) Small model network consisting of 8 bilateral neuron pairs. Dashed connectors represent paired synapses. Continuous connectors are unpaired synapses. Darkened connectors correspond to the examples in the text. (B) Fraction of unpaired synapses, 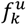, in the different connectome networks. Shown also is the average (black bar), *f*^*u*^, for each network. (C) Comparison of 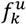values between connectome networks. *ρ* - Spearman rank correlation. n=83, p<0.0001 for all plots. (D) Comparison between 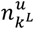and 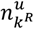, the number of unpaired synapses of the left vs. right member of each neuron class, for each connectome network. *ρ* - Spearman rank correlation. n=83 for all plots. p=0.0004 for CeH1c and p<0.0001 for CeH2c and CeH3c networks.

In general, we define the fraction of unpaired synapses of neuron class *k* as 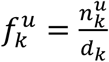, where 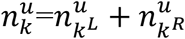is the sum of unpaired synapses of the left and right members of neuron class *k*. An 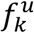value close to 0, indicates a neuron class with high symmetry in synaptic composition. Values close to 1 represent high asymmetry in synaptic composition. Computing the fraction of unpaired synapses, 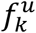, in the different connectome networks revealed a large spread of values, with comparable averages, 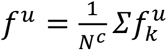(Fig. 2B), where *N*^*c*^ is the number of neuron classes. Not even one neuron class in any network was perfectly symmetrical 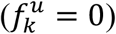. 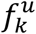values showed a significant correlation between networks (Fig. 2C), suggesting that the extent of asymmetry in synaptic composition of each neuron class is a conserved feature of that neuron class. We also found a significant correlation between 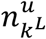and 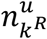in all networks (Fig. 2D). Taken together, our results show that bilateral members of a neuron class exhibit asymmetry in the composition of a subset of their synapses, and this subset shows similarity in size in the two neurons.

### Stochasticity contributes to asymmetry in synaptic composition

Asymmetry in synaptic composition (Fig. 2) could stem from predetermined genetic^23^ or biomechanical^24^ factors, or from ongoing adjustments by activity-dependent^25^ or stochastic^26^ processes. To distinguish between these possibilities, we examined the consistency of all chemical synaptic connections across the 3 connectome networks (Fig. 3A). We reasoned that consistent synapses are more likely to be predetermined than adjusted. The average portion of all chemical synapses present in all 3 networks was surprisingly low, ∼24% (Fig. 3A, left). Out of these, the share of consistent paired synapses was significantly higher, ∼39% (Fig. 3A, middle; Z=9.51, p<0.00001). In contrast, the average portion of unpaired synapses included in all networks was only ∼2.5% (Fig. 3A, right), significantly smaller than the fraction of paired synapses (Z=19.67, p<0.00001). This finding suggests that unpaired synapses are particularly variable across individuals, and are therefore more likely to be determined by stochastic or experience-dependent processes than paired synapses.

**Figure 3.**
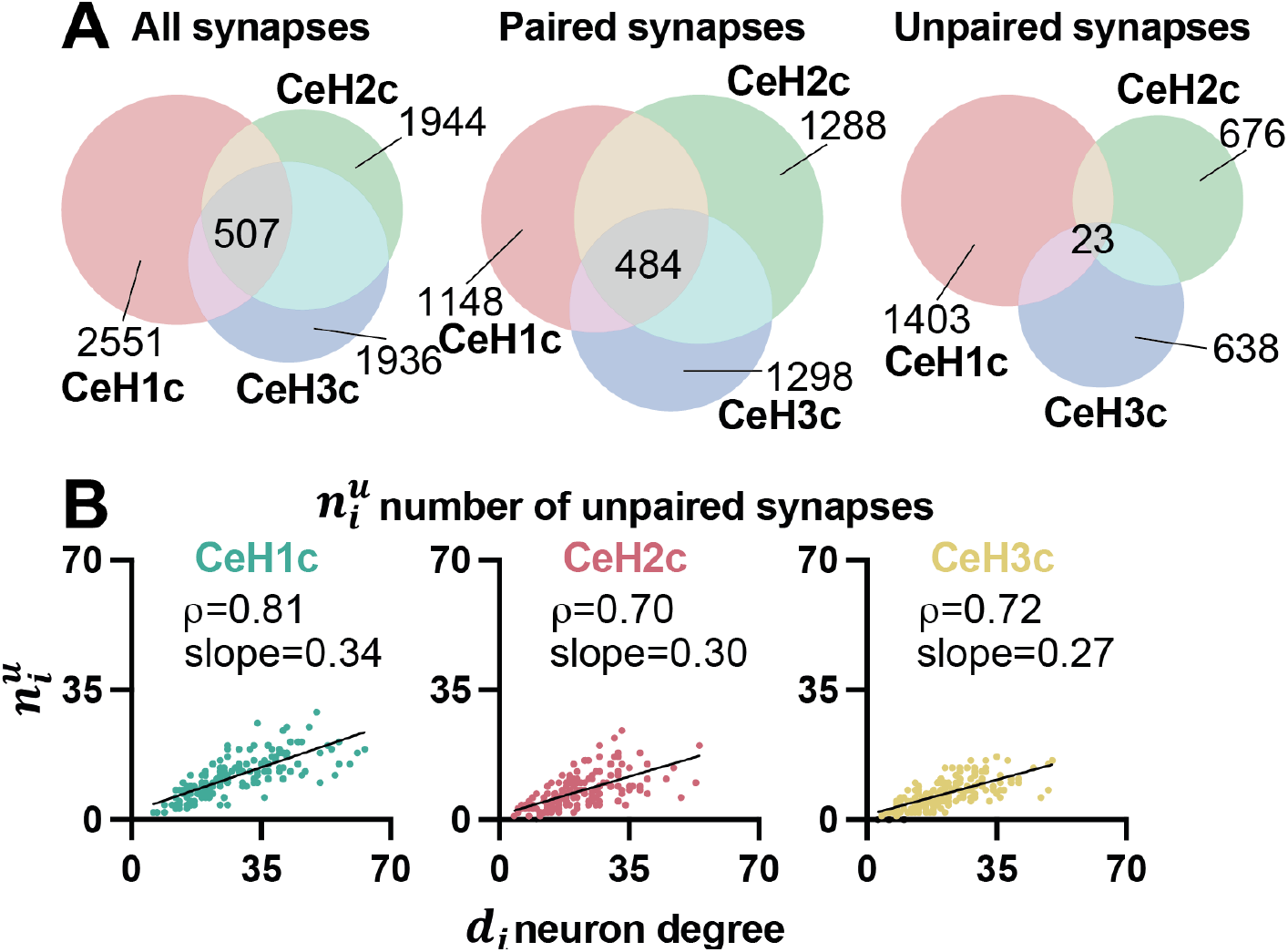
Asymmetry in chemical synaptic composition is stochastically driven. (A) Venn diagram showing the number of all (left), paired (middle) and unpaired (right) chemical synapses in each connectome network and the number of such synapses shared by all three networks (center of each diagram). (B) Comparison between 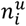, the number of unpaired synapses of each individual neuron,and *d*_*i*_, the neuron’s degree for each connectome network. *ρ* - Spearman rank correlation. Slopes fitted from simple linear regression. n=166, p<0.0001 for all plots.

To extend our analysis, we examined the relation between 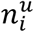, the number of unpaired synapses of each individual neuron, and *d*_*i*_, the neuron’s degree. These two variables showed significant correlation in all networks (Fig. 3B), indicating that the number of unpaired synapses of each neuron is proportional to the neuron’s degree. This proportion can be estimated from the slopes in Fig. 3B, whose values across networks were quite similar, suggesting a consistent fixed fraction of unpaired synapses in *C. elegans* neurons. These results point at a possible stochastic process underlying the choice of unpaired synapses, presumably tuned by a global fixed probability for the formation of unpaired synapses in each neuron, as opposed to an experience-dependent process, expected to show much more heterogeneity across neurons (since plasticity is presumably selective, affecting only specific neurons and synapses^27^). Taken together, our analysis suggests that within each individual neuron, a fixed portion of unpaired synapses is formed through a stationary stochastic process.

### Symmetry in synapse composition has a non-stochastic component

As we have found, paired (symmetrical) synapses appear to be more consistent than unpaired synapses across connectome networks (Fig. 3A), suggesting that they are less driven by stochasticity than unpaired synapses. We sought to study the extent to which randomness may nevertheless guide symmetry in synaptic composition through stochastic formation of paired synapses. To this end we considered network directionality, the arrangement of chemical synapses so that transmission occurs only in one direction (from presynaptic to postsynaptic neuron).

A completely random directed network is expected to contain more unpaired synapses than an equivalent undirected network, maintaining all synaptic connections, but ignoring their directionality. This is because the directed network offers more opportunities for unpaired synapses to occur. For example, in the undirected small model network (Fig. 4A, left), the AL-BL synapse has a contralateral counterpart, AR-BR, and is therefore considered a paired synapse (dashed connector). If directionality is added to the network, then an AL→BL connection, for instance (Fig. 4A, right), now has a 50% chance of being unpaired instead of paired (AL→BL would pair with AR→BR, but not with BR→AR), reducing the relative number of paired connections and increasing asymmetry. To illustrate, we generated a series of small random undirected networks containing *N* = 40 neurons and *d* = 80 synapses and symmetrized or desymmetrized them (see below) so that their average fraction of unpaired synapses, *f*^*u*^, was 0.00, 0.34 or 0.84. We derived corresponding directed versions of these networks, by assigning random directionality to all their synapses. In order to assess overall network asymmetry, we computed *f*^*u*^ for each undirected and equivalent directed network. As we had postulated for such stochastic networks, *f*^*u*^ was substantially higher in the directed networks compared to the undirected ones (Fig. 4Bi), with the gap between undirected and directed *f*^*u*^ decreasing as overall network asymmetry increased (Fig. 4Bii). The average 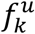correlation between undirected and directed networks (with mean undirected *f*^*u*^ = 0.34) was moderate (Fig. 4Biii), and the average slope of the linear regressions constrained by these values was considerably smaller than 1 (Fig. 4Biii), consistent with greater asymmetry of directed compared to undirected random networks.

**Figure 4.**
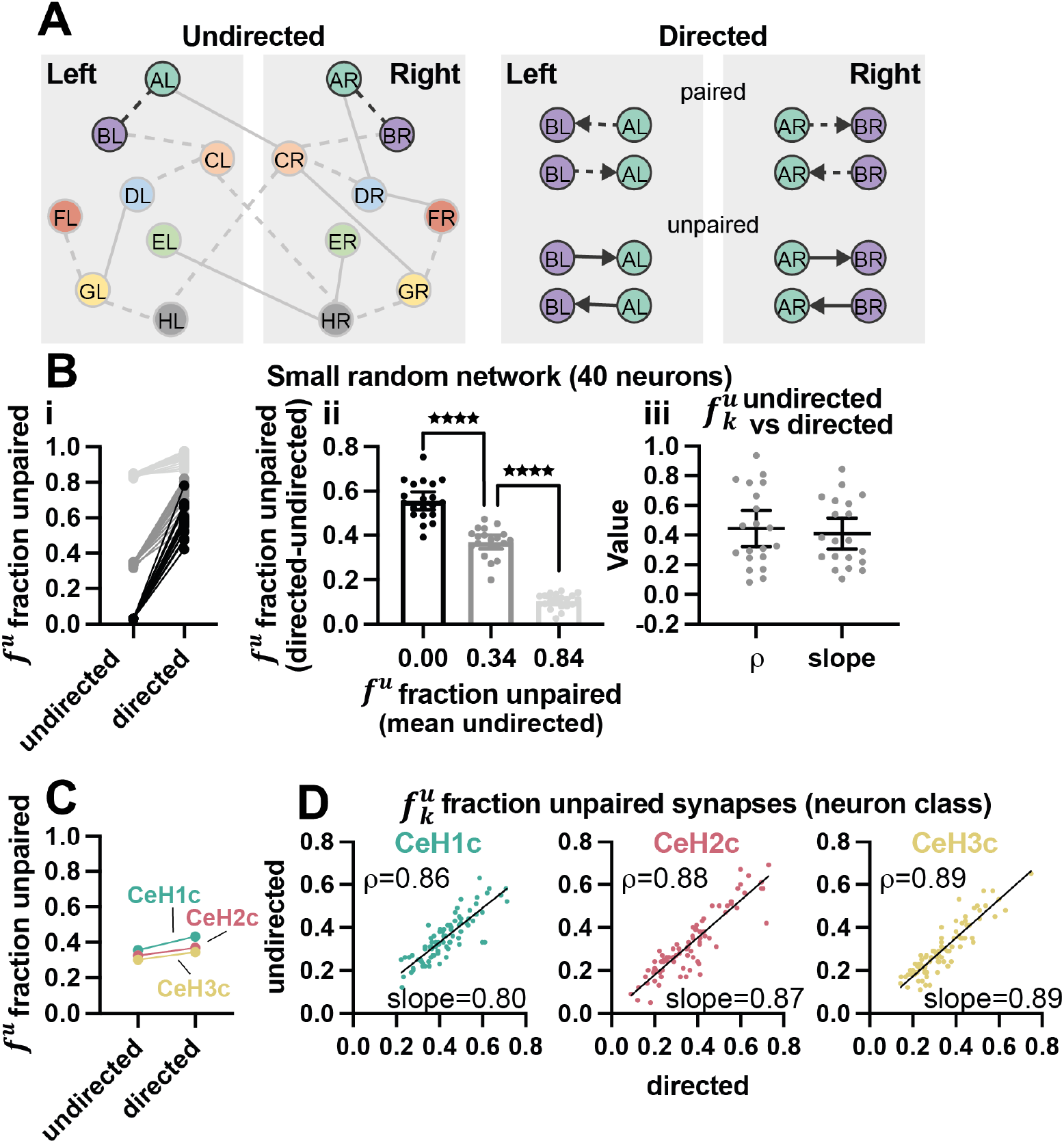
Symmetry in chemical synaptic composition is not strongly stochastic. (A) Left, small undirected model network consisting of 8 bilateral neuron pairs. Dashed connectors represent paired synapses. Continuous connectors are unpaired synapses. Darkened connectors point to an example connection between the A and B neuron classes. Right, several possible directed connections between the A and B neuron classes. The top two options represent paired connections, and the bottom two unpaired connections. (B) Small undirected and matched directed random networks consisting of 40 neurons and 80 interconnecting synapses grouped according to their average fraction of undirected unpaired synapses, *f*^*u*^ = 0.00, 0.34, 0.84. (i) *f*^*u*^ compared between undirected and directed versions of each network. (ii) Difference in *f*^*u*^ between directed and undirected versions of each network. n=20. One-way ANOVA p<0.0001. Post hoc Dunnett′s multiple comparisons test, p<0.0001 for both comparisons. (iii) Results of comparison of 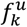between undirected and directed versions of each network (with average undirected *f*^*u*^ = 0.34). Shown are the Spearman rank correlation coefficient, *ρ*, values and the slope as fitted by a linear regression, for each network instance. Error bars represent 95% confidence intervals. Each dot represents a different network. (C) Fraction of unpaired synapses, *f*^*u*^, in directed vs. undirected connectome networks. (D) Comparison of the fraction of neuron class unpaired synapses, 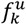, between undirected and directed connectome networks. *ρ* - Spearman rank correlation coefficient. Slope as fitted by a linear regression. n=83, p<0.0001 for all plots.

In order to assess stochasticity in the specification of *C. elegans* paired synapses, we thus compared *f*^*u*^ values between the original directed connectome networks and corresponding networks, constructed to be undirected. A large decrease in *f*^*u*^ (as in the purely random networks; Fig. 4B) would indicate a strong stochastic influence on the emergence of paired synapses. In reality, the calculated differences in *f*^*u*^ values between directed and undirected connectome networks were rather minor (compare Fig. 4C to Fig. 4Bi), and the fraction of unpaired synapses within each neuron class, 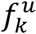(Fig. 4D), and within each individual neuron, 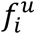(Fig. S2), showed strong correlations between the undirected and directed connectomes with slopes close to 1 (compare with Fig. 4Biii). These results provide further evidence for regulated rather than random specification of symmetrical synapses, or at least, tight coordination in synaptic choice between neuron class members.

### Redundancy and reachability reflect a tradeoff between network symmetry and asymmetry

The *f*^*u*^ values calculated from the *C. elegans* chemical connectomes reveal a considerable level of asymmetry in synaptic connectivity (Fig. 4C). What are the functional implications of such asymmetry? To begin to address this question we sought to identify specific network properties that capture functional differences between symmetry and asymmetry. To gain a better grasp of the issue, we turn once again to small example networks, this time ranging from fully symmetrical (Fig. 5A; *f*^*u*^ = 0.0) to fully asymmetrical (Fig. 5C; *f*^*u*^ = 1.0). We accompany each detailed individual neuron network with a collapsed version of that network, showing the connections between its neuron classes (Fig 5A-C). We assume that the members of each neuron class perform similar or related functions. We note that bilateral members of half (53.6%) of all neuron classes are interconnected by at least one chemical or electrical synapse (in the most detailed connectome network, CeH0), potentially coupling their activity. As can be seen, the same number of synapses are distributed very differently in symmetrical vs. asymmetrical networks. In the symmetrical network (Fig. 5A, right) fewer classes are connected to each other compared to the asymmetrical network (Fig. 5C, right). However, each inter-class connection is stronger, comprising more synapses (Fig. 5A, right). This tradeoff in synapse distribution may reflect an important balance between symmetrical and asymmetrical network arrangements. We frame this tradeoff in terms of Redundancy vs. Reachability.

**Figure 5.**
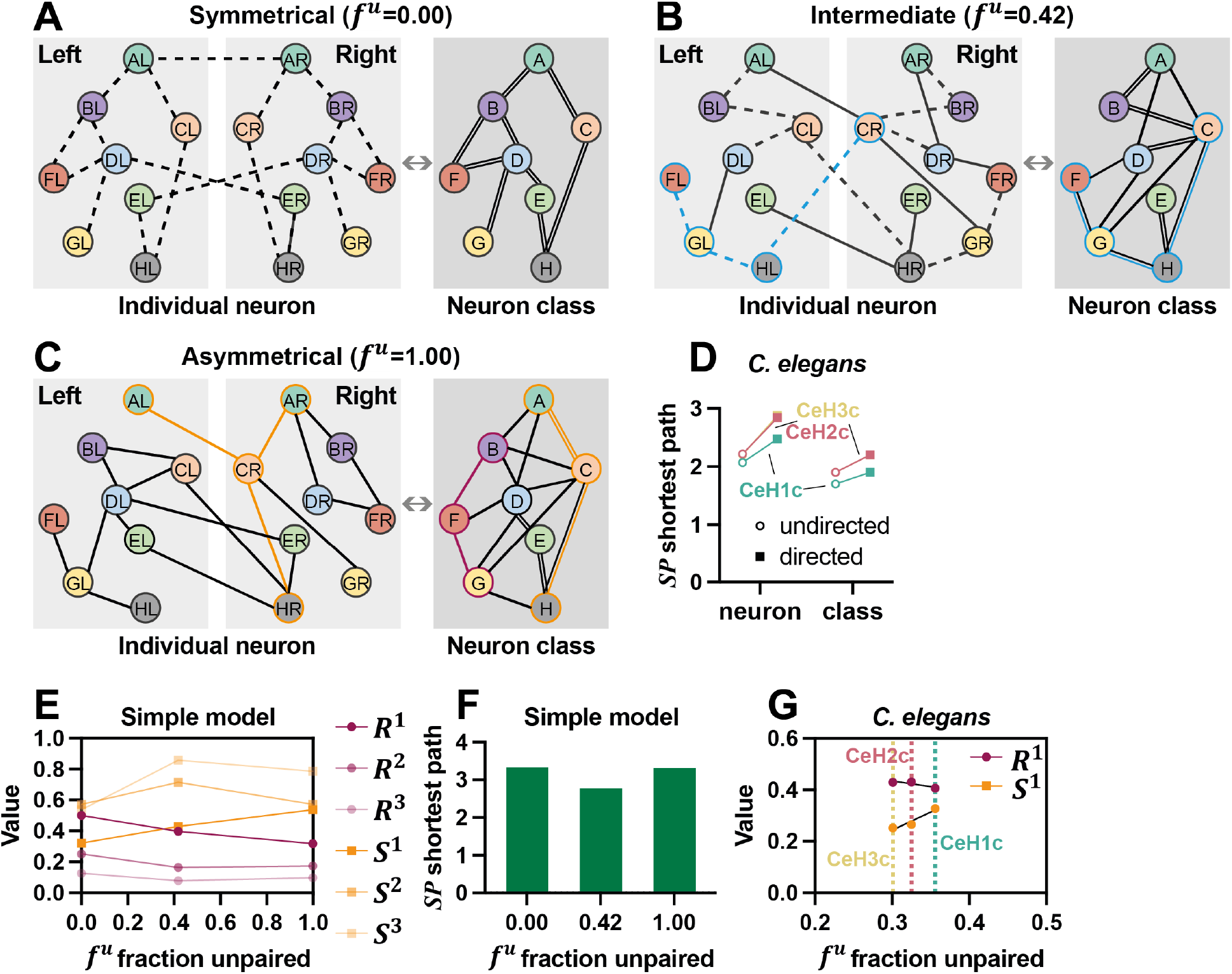
Redundancy and Reachability in a small model network and in the *C. elegans* connectomes. (A-C) Small model networks consisting of 8 bilateral neuron pairs interconnected by 19 undirected synapses. Dashed connectors represent paired synapses. Continuous connectors are unpaired synapses. Each network is presented at the individual neuron level (left) and at the neuron class level (right). The number of connectors between each two neuron classes indicates the number of synapses connecting between their individual neuron members. The networks range from (A) fully symmetrical (*f*^*u*^ = 0.00), through (B) intermediate (*f*^*u*^ = 0.42), to (C) fully asymmetrical (*f*^*u*^ = 1.00). (D) Mean shortest path length (*SP*) connecting any two neurons (left) or any two neuron classes (right) in the various *C. elegans* connectome networks. n=180 individual neurons, n=83 neuron classes of bilateral neuron pairs, and n=14 neuron classes of single non-bilateral neurons. (E) Redundancy, *R*^*n*^, and Reachability, *S*^*n*^, plotted against the fraction of unpaired synapses, *f*^*u*^, of the three model networks (A-C). (F) Mean shortest path length (*SP*) connecting any two neurons in the small model networks (A-C). (G) Redundancy, *R*^1^, and Reachability, *S*^1^, plotted against the fraction of unpaired synapses, *f*^*u*^, of the undirected *C. elegans* connectome networks. Dotted lines indicate the *f*^*u*^ of each network.

We refer to *Redundancy* as an abundance of synaptic connections linking specific neuron classes. Redundancy could selectively enhance or refine particular routes of information flow in the circuit, and when necessary, serve as back up in response, for example, to weakened signaling or neural damage. For a given number of synapses in a network, greater symmetry in synaptic composition should lead to increased Redundancy (owing to duplication).

We denote by *Reachability* the number of distinct neuron classes that can be reached by synaptic connections from each neuron class. Since asymmetrical networks include more diverse synaptic connections than symmetric networks (at the expense of duplication), increased asymmetry should raise the number of interconnected neuron classes, thus enhancing Reachability.

### Redundancy and Reachability vary as a function of network symmetry vs. asymmetry

In order to formally characterize Redundancy and Reachability, we first introduce several useful terms. In a given network, a *path* of length *n* is a sequence of *n* + 1 distinct neurons connected by *n* synapses. For example, in Fig. 5B (left), CR-HL-GL-FL (blue path) is a path of length *n* = 3 comprising 4 neurons. We distinguish between a *neuron path*, composed of a sequence of individual neurons, and a *class path*, composed of a sequence of neuron classes (disregarding the identity of individual neuron members of each class), in the example, C-H-G-F is a class path (Fig. 5B, right).

We define a *realizable* class path as a class path, for which at least one corresponding neuron path exists in the network. For example, in Fig. 5C (right), the class path B-F-G (purple path) is not realizable, since there is no corresponding neuron path (Fig. 5C, left) that can instantiate this class path. In contrast, A-C-H (orange path) is realizable by two neuron paths: AL-CR-HR and AR-CR-HR. The theoretical maximal number of neuron paths that could instantiate a particular class path, *j*, of length *n* in an undirected (see below) network is 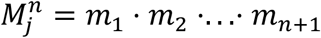, where *m*_*k*_ denotes the number of neuron members belonging to neuron class *k* (in the case where all neuron classes in class path *j* are bilateral neuron pairs, then 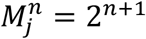). Since neural networks are rarely, if ever, complete (*i*.*e*., including synapses between every possible pair of neurons), the actual number of neuron paths, 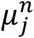, that realize a specific class path, *j*, of length *n* should be typically much smaller than the maximal possible value 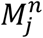. For example, as shown above, the 2 realizable neuron paths for class path A-C-H are few relative to 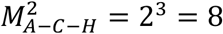. Notably, many class paths lack any corresponding neuron paths whatsoever, and thus are not realizable (*e*.*g*., class path B-F-G in Fig. 5C, described above).

Thus, the evaluation of network Redundancy and Reachability relies on path analysis. This may be considerably simplified if performed in undirected rather than directed networks. Such an approximation can be justified by the similarities we have found between undirected and directed *C. elegans* connectome networks with respect to symmetry and asymmetry (Fig. 4C,D). In addition, we find the mean shortest path length (*SP*) connecting any two neurons, and especially any two neuron classes, to be comparable between directed and undirected *C. elegans* connectome networks (Fig. 5D), and to covary within each network (*e*.*g*., a network with smaller undirected *SP* shows also smaller directed *SP* compared to other networks). Thus, the removal of directionality constraints may result in quantitative changes to certain network properties. However, the qualitative effects of this approximation on network paths seem to be minor.

We formally define network Redundancy, 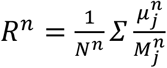, as the average number, 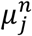, of neuron paths of length *n* that realize each class path, *j*, in the network, where *N*^*n*^ is the number of realizable class paths of length *n*. Importantly, only realizable class paths are included in the calculation. In addition, 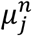is normalized (before averaging) by the maximal possible number of neuron paths, 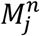, that could constitute the class path. *R*^*n*^ may thus range between 0 and 1. However, since only realizable class paths are considered, and the network contains a minimal number of synaptic connections, the actual minimal *R*^*n*^ value will be higher than 0. To illustrate the relations between *R*^*n*^ and network symmetry vs. asymmetry, we derive *R*^1^ for the model networks in Fig. 5. In the fully symmetrical network (Fig. 5A; *f*^*u*^ = 0) all realizable class paths of length *n* = 1 have 2 corresponding neuron paths (double connectors in Fig. 5A, right) out of a maximum of 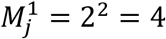. Thus, for this network, *R*^1^ = 0.50 (Fig. 5E). As asymmetry increases (Fig. 5B; *f*^*u*^ = 0.42) Redundancy decreases, *R*^1^ = 0.40 (Fig. 5E), and is lowest, *R*^1^ = 0.32 for the fully asymmetric configuration (Fig. 5C,E; *f*^*u*^ = 1). For longer path lengths of *n* > 1, overall *R*^*n*>1^ levels drop, and *R*^*n*^ compared to *R*^1^ shows decreased dependence on *f*^*u*^ (Fig. 5E). This is because 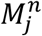grows exponentially with *n*, whereas the extent of duplication doesn’t change. It is therefore sufficiently informative to use *R*^1^ as a measure of network redundancy without the need to consider longer path lengths.

We define network Reachability, 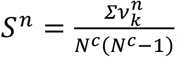, as the average number, 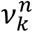, of neuron classes that can be reached from each neuron class, *k*, by a realizable class path of length, *n. N*^*c*^ is the number of all neuron classes. 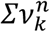is normalized by the maximal possible number of classes that could be reached from each neuron class, *N*^*c*^− 1. The values of *S*^*n*^ may range from 0 to 1, but are in practice, higher than 0, since it can be assumed that each neuron in the network has at least one synaptic contact, enabling minimal reachability. To demonstrate how *S*^*n*^ relates to network symmetry vs. asymmetry we compute *S*^1^ for the model networks in Fig. 5. In the symmetrical network (Fig. 5A; *f*^*u*^ = 0.00), the average number of neuron classes reachable from each neuron class via a realizable class path of length *n* = 1 is *S*^1^ = 0.32 (Fig. 5E). As asymmetry increases (Fig. 5B; *f*^*u*^ = 0.42) so does Reachability, *S*^1^ = 0.43 (Fig. 5E), and is highest, *S*^1^ = 0.54, for the fully asymmetric network (Fig. 5C,E; *f*^*u*^ = 1.00). For paths of length *n* > 1, *S*^*n*^ values generally rise (Fig. 5E). This is because the average shortest path length of each of these networks is close to 3 (Fig. 5F), so that paths of length *n* = 2 already enable reaching many neuron classes. Indeed, the relatively lower *SP* of the intermediate network (Fig. 5F; *f*^*u*^ = 0.42) shows also higher *S*^2^ and *S*^3^, compared to the fully symmetrical (*f*^*u*^ = 0.0) or fully asymmetrical (*f*^*u*^ = 1.0) networks. Therefore, similarly to *R*^1^, we find also *S*^1^ to be sufficient for capturing network reachability in a straightforward manner.

The *C. elegans* connectome is obviously more complex than a small model network, differing from it in topology and other aspects. We calculated *R*^1^ and *S*^1^ for the undirected connectome networks (Fig. 5G). Notably, even within the small range of *f*^*u*^ values of the different networks, *R*^1^ covaried negatively, and *S*^1^ covaried positively with *f*^*u*^ (Fig. 5G), similar to the model network (Fig. 5E). These results demonstrate the sensitivity and applicability of Redundancy and Reachability in the analysis of connectome symmetry vs. asymmetry.

### The level of asymmetry in the connectome may favor Redundancy over Reachability

What could the *f*^*u*^ values of the *C. elegans* chemical synaptic connectome tell about the balance in these networks between Redundancy and Reachability? To address this question, it is useful to first appreciate how *R*^1^ and *S*^1^ are related to *f*^*u*^, and how sensitive these measures are to differences in *f*^*u*^ values. A linear dependency, for example (Fig. 6A), would imply that smaller *f*^*u*^ values, as occurs in the connectome (Fig. 4C, Fig. 5G) should correspond to a preference for Redundancy at the expense of Reachability. In this (linear) case, the tradeoff between *R*^1^ and *S*^1^ will depend also on their respective slopes. For instance, a steep *R*^1^ slope in conjunction with a shallow *S*^1^ slope would indicate that Redundancy (*R*^1^) can be gained or lost with little impact on Reachability (*S*^1^). Alternatively, a non-linear relationship between *R*^1^ or *S*^1^ and *f*^*u*^ (*e*.*g*., Fig. 6B) may entail other preferences and tradeoffs balanced by the level of network symmetry vs. asymmetry.

**Figure 6.**
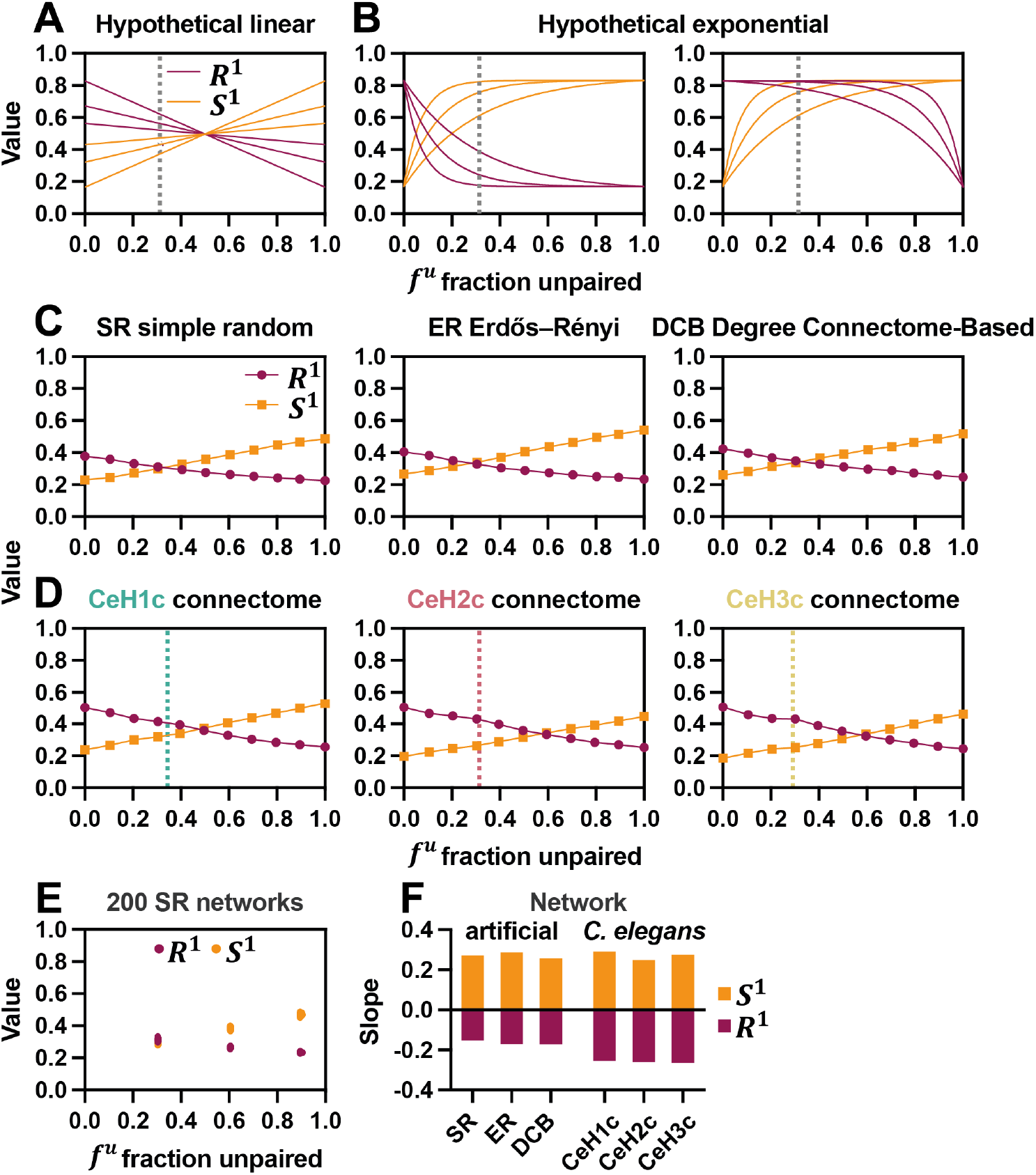
A symmetrical bias of the connectomes favors Redundancy over Reachability. (A) Hypothetical linear dependency between *R*^1^ and *S*^1^, and *f*^*u*^ with varying slopes. (B) Hypothetical exponential dependency between *R*^1^ and *S*^1^, and *f*^*u*^ with varying exponents. (C) Redundancy, *R*^1^, and Reachability, *S*^1^, plotted against the fraction of unpaired synapses, *f*^*u*^, for various large artificial random networks designed to vary in *f*^*u*^ from 0.0 to 1.0. (D) Redundancy, *R*^1^, and Reachability, *S*^1^, plotted against the fraction of unpaired synapses, *f*^*u*^, for the three *C. elegans* connectome networks systematically modified to vary in *f*^*u*^ from 0.0 to 1.0. Dotted lines indicate actual *f*^*u*^ value for each network. (E) Redundancy, *R*^1^, and Reachability, *S*^1^, for 3 sets of 200 simple random (SR) networks with 3 different levels of asymmetry (*f*^*u*^ = 0.3, 0.6, 0.9). (F) Slopes of the *R*^1^ and *S*^1^ vs. *f*^*u*^ plots from (C,D) as fitted by linear regression.

To determine how *R*^1^ and *S*^1^ are governed by *f*^*u*^, we generated several artificial networks similar in dimension to the connectome networks (see Methods): SR, a simple random network. ER, an Erdős–Rényi random network^28,29^, and DCB a random network with initial neuron degree values, *d*_*i*_, and a restricted synaptic partner pool based on the connectome and ‘contactome’ (list of all neuronal adjacencies^30,31^; see Methods). We paired the neurons in the networks to form bilateral neuron classes, as in the connectome. We devised an algorithm (see Methods) for symmetrizing or desymmetrizing the network, based on iterative replacement of random synaptic connections, to obtain a series of networks with *f*^*u*^, ranging from complete symmetry (*f*^*u*^ = 0) to full asymmetry (*f*^*u*^ = 1). We computed for each such series of networks *R*^1^ and *S*^1^ at fixed *f*^*u*^ steps. Plotting *R*^1^ and *S*^1^ against *f*^*u*^ (Fig. 6C) revealed a highly linear dependency of both *R*^1^ and *S*^1^ on *f*^*u*^ (Fig. 6C). We found a similar linear relationship also when we subjected the original connectome networks to symmetrization and desymmetrization (Fig. 6D). Such linearity is suggested also by the comparison of *R*^1^ and *S*^1^ between the 3 connectome networks (Fig. 5G).

To further examine the strength of the association between *R*^1^, *S*^1^ and *f*^*u*^, We generated 200 random instances of the SR network, which were symmetrized or desymmetrized to obtain networks with particular *f*^*u*^. The resulting *R*^1^ and *S*^1^ values were restricted to a considerably narrow range (Fig. 6E), emphasizing the tight relations between Redundancy, Reachability and network symmetry vs. asymmetry.

We performed linear regression to each of the *R*^1^ and *S*^1^ curves and attained the fitted slopes (Fig. 6F). Whereas the *R*^1^ slopes were similar in magnitude to *S*^1^ in the connectome-generated networks (Fig. 6D,F), the artificial networks showed smaller *R*^1^ slopes compared to *S*^1^ (Fig. 6C,F). This difference could stem from particular topological features distinctive of the real connectome networks. The similar slopes of *R*^1^ and *S*^1^ (Fig. 6D,F) position the putative crossover point between Redundancy and Reachability in the middle between symmetry and asymmetry (*f*^*u*^ ≈ 0.5). Since we find *C. elegans f*^*u*^ values to be somewhat biased towards 0 (Fig. 6D, dotted lines), this suggests that the layout of its connectome prioritizes Redundancy over Reachability, so that relatively fewer but stronger inter-class connections are preferred over broader but weaker connectivity.

### Paired and unpaired synapses play distinct role in tuning Redundancy and Reachability

As we have shown, paired synapses inherently reinforce specific class paths through duplication, thus promoting Redundancy. However, unpaired synapses may also contribute to Redundancy. For example (Fig. 5B), the two unpaired synapses, EL-HR and ER-HR, provide two alternative connections for realizing the E-H class path, and thus contribute to Redundancy despite being unpaired. In the extreme case, of a fully asymmetrical network (*f*^*u*^ = 1), all synapses are unpaired, and so any Redundancy in such a network is exclusively due to these unpaired connections. To directly probe the particular contribution of paired synapses to Redundancy, we calculated *R*^*p*,1^, a metric similar to *R*^1^, but comprising only paired synapses (with the average, however, taken for all realizable class paths, as in *R*^1^, regardless of whether they include paired or unpaired synapses; see Methods). We plotted *R*^*p*,1^ for networks of different *f*^*u*^ values for the three connectomes (Fig. 7A). As expected, in fully symmetrical (*f*^*u*^ = 0) networks, *R*^*p*,1^ = *R*^1^, since Redundancy in this case is entirely established by paired synapses. In contrast, *R*^*p*,1^ equaled 0 for fully asymmetrical (*f*^*u*^ = 1) networks, since Redundancy is entirely due to unpaired synapses. Calculating the fraction of *R*^*p*,1^ out of the total *R*^1^, 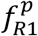, for the actual connectome *f*^*u*^ values (Fig. 7A, dotted lines), revealed that paired synapses contribute a substantial portion of Redundancy to the network (∼65%), but unpaired synapses participate as well (∼35%; Fig. 7B).

**Figure 7.**
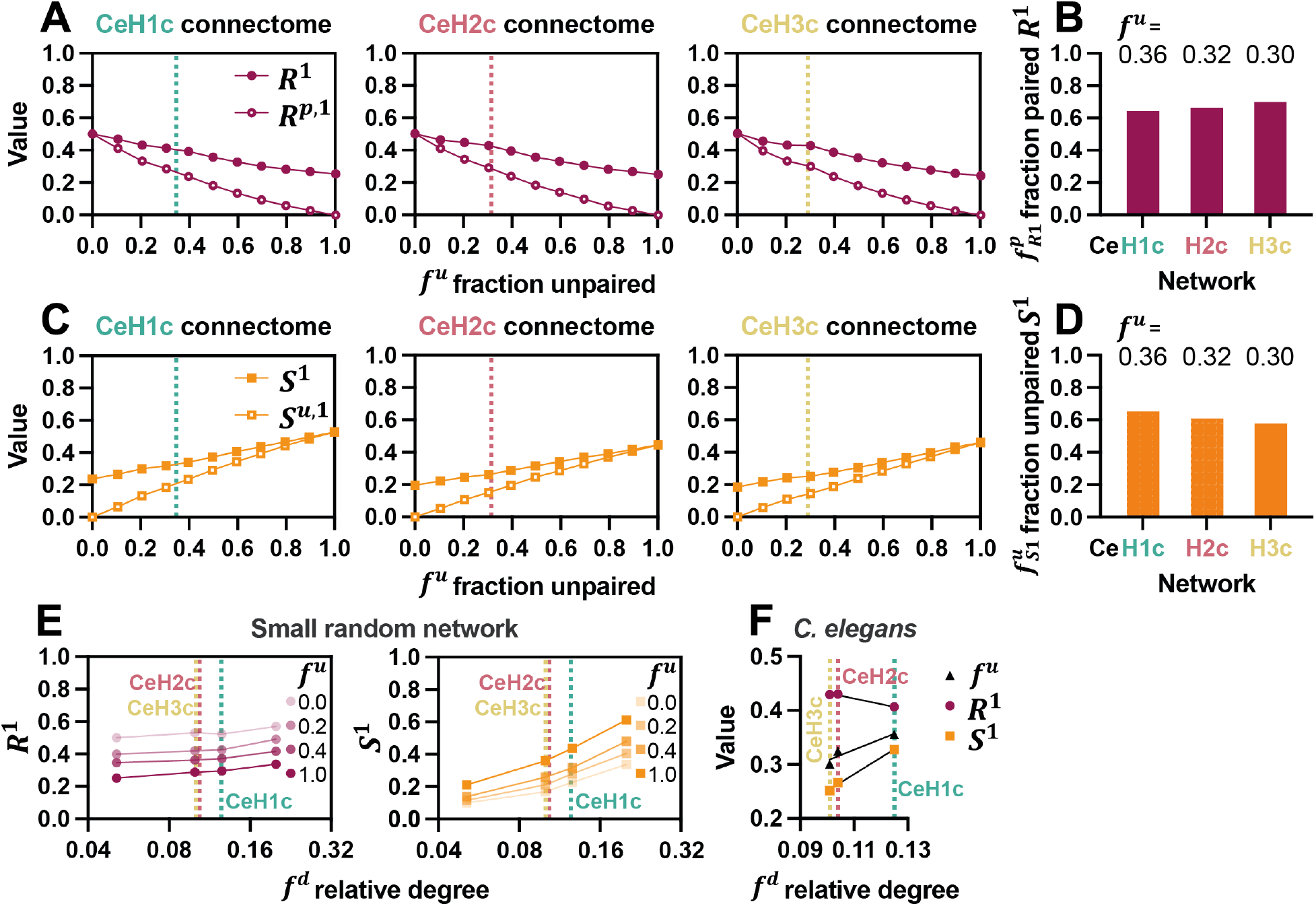
The influence of relative network degree on Redundancy and Reachability. (A) Redundancy due exclusively to paired synapses, *R*^*p*,1^, plotted against the fraction of unpaired synapses, *f*^*u*^, for the three *C. elegans* connectome networks systematically modified to vary in *f*^*u*^ from 0.0 to 1.0. Dotted lines indicate actual *f*^*u*^ value for each network. (B) 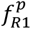, fraction of Redundancy due only to paired synapses, *R*^*p*,1^, out of total Redundancy, *R*^1^. (C) Reachability due exclusively to unpaired synapses, *S*^*u*,1^, plotted against the fraction of unpaired synapses, *f*^*u*^, for the three *C. elegans* connectome networks systematically modified to vary in *f*^*u*^ from 0.0 to 1.0. Dotted lines indicate actual *f*^*u*^ value for each network. (D) 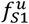, fraction of Reachability due to unpaired synapses, *S*^*p*,1^, out of total Reachability, *S*^1^. (E) Redundancy, *R*^1^, and Reachability, *S*^1^, of a series of small random networks (40 neurons) with varying number of synapses (*f*^*d*^ = 0.05, 0.1, 0.125, 0.2) and fraction of unpaired synapses, *f*^*u*^. Dotted lines indicated *C. elegans* connectome networks *f*^*d*^ values. (F) Redundancy, *R*^1^, Reachability, *S*^1^, and fraction of unpaired synapses, *f*^*u*^, plotted against *C. elegans* connectome networks *f*^*d*^ values.

We also evaluated the relative contribution of unpaired synapses to Reachability. To this end we calculated *S*^*u*,1^ similarly to *S*^1^, considering for each neuron class only connected partner classes that are linked via unpaired synapses (Fig. 7C). Obviously, for fully symmetrical networks, comprising exclusively paired synapses (*f*^*u*^ = 0), *S*^*u*,1^ = 0, and in fully asymmetrical networks lacking any paired synapses (*f*^*u*^ = 1), *S*^*u*,1^ equals *S*^1^. The ratio of *S*^*u*,1^ to *S*^1^, 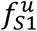, for the connectome *f*^*u*^ values was rather high (average ∼60%; Fig. 7D), indicating that Reachability is largely based on unpaired synapses, but also paired synapses have an appreciable share (∼40%).

### Network degree affects the dependency of Redundancy and Reachability on network symmetry

Network degree, *d* (the total number of synapses in the network), may affect the relations between asymmetry, Redundancy and Reachability. To examine this, we studied the effects of varying 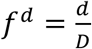, the ratio between *d* and 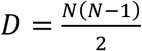(the maximal possible degree of a complete network, where all neurons are connected to all other neurons) on *R*^1^ and *S*^1^. On the one hand, extremely sparse networks with low *f*^*d*^ are likely to be inherently asymmetrical since they present less opportunities for synaptic pairing. This would lead to low Redundancy, but also low Reachability and moderate dependency on *f*^*u*^. The opposite is true for very dense networks, which should be distinctly symmetrical with high, close to constant, Redundancy and Reachability. To examine intermediate *f*^*d*^ values, we constructed a series of small (40 neuron) undirected simple random networks with variable *f*^*d*^ (Fig. 7E; Fig. S3), including the *f*^*d*^ values of the *C. elegans* connectome networks (Fig. 7E, dashed lines; Fig. 7F). Interestingly, within this range, *R*^1^ showed weak dependency on *f*^*d*^, compared to *S*^1^, which increased considerably with *f*^*d*^, for a broad range of *f*^*u*^ values (Fig. 7E). This difference between seemingly scale-independent *R*^1^ and scale-dependent *S*^1^ stems from two distinct effects of network degree within intermediate values. First, although a relatively larger *f*^*d*^, implying more synapses, produces more realizable class paths, the number of alternative neuron paths that realize these paths remains similar, unaffecting overall Redundancy. Second, a denser network increases the possibilities for each neuron class to connect to other classes, thus enhancing Reachability. Therefore, in theory, in the *C. elegans* connectome, Reachability (*S*^1^) could have been larger without requiring an increase in asymmetry (*f*^*u*^) or a decrease in Redundancy (*R*^1^), if the total number of synapses were larger. In reality, it appears that a smaller network degree is favored over increased Reachability.

### Different connectomes vary mostly in asymmetry and Reachability

To conclude our investigation, we wished to expand our analysis and probe additional available connectome networks (Fig. 8A). First, we examined the full corrected original *C. elegans* connectome, CeH0c, from which we had derived CeH1c in the preceding analyses^20^. We found CeH0c to be comparable to the other networks that we have used (Fig. 8B), and used this network for further analyses. Overall comparison between various connectomes (shaded connectomes in Fig. 8A) revealed a relatively broad distribution of symmetry vs. asymmetry (Fig. 8C; *f*^*u*^) and Reachability (Fig. 8C; *S*^1^). Redundancy, in contrast, showed a much narrower range of values (Fig. 8C; *R*^1^), in accordance with our finding that Redundancy is less sensitive to network configuration than Reachability (Fig. 7E). The average contribution of paired synapses to Redundancy and unpaired synapses to Reachability was similar across connectomes (Fig. 8C; 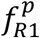and 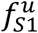, respectively).

**Figure 8.**
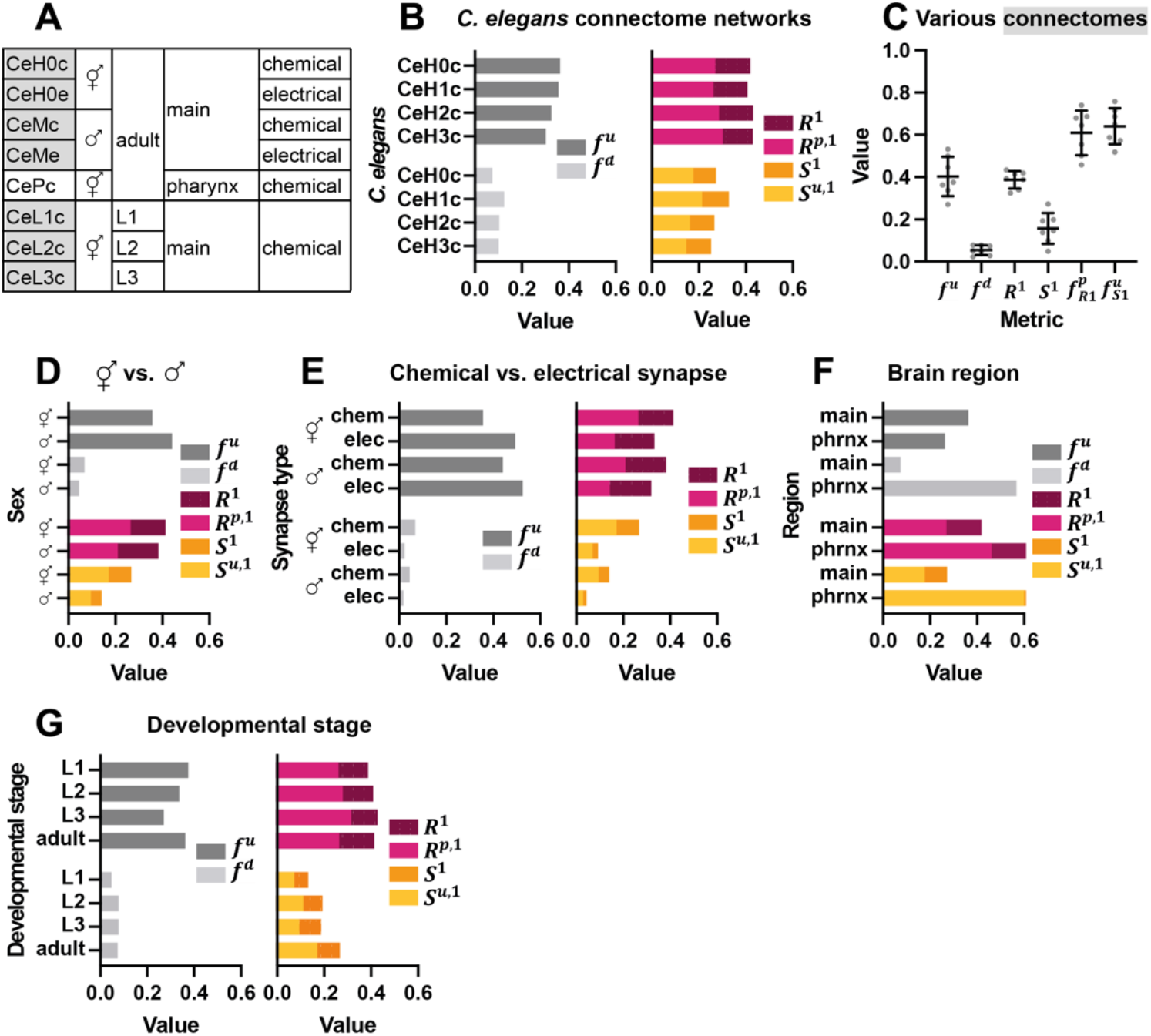
Different connectomes vary mostly in asymmetry and Reachability and less so in Redundancy. (A) List of additional *C. elegans* connectomes analyzed from different sexes, brain region, type of synapse (chemical vs. electrical) and developmental stage. (B) Comparison between the complete *C. elegans* chemical connectome network, CeH0c, and the connectome networks used throughout the study, CeH1c, CeH2c, CeH3c. (C) Distribution of *f*^*u*^, *f*^*d*^, *R*^1^, *S*^1^, 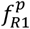and 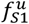, for the connectome networks shaded in (A). Error bars indicate mean ±standard deviation. (D) Comparison of asymmetry, Redundancy and Reachability between hermaphrodite (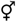; CeH0c) and male (♂; CeMc) chemical connectome networks. (E) Comparison of asymmetry, Redundancy and Reachability between chemical (CeH0c and CeMc) and electrical (CeH0e and CeMe) connectome networks. (F) Comparison of asymmetry, Redundancy and Reachability between main (CeH0c) and pharynx (CePc) chemical connectome networks. (G) Comparison of asymmetry, Redundancy and Reachability between different developmental (CeL1c, CeL2c, CeL3c) and adult (CeH0c) chemical connectome networks.

Comparing hermaphrodite (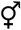; CeH0c) and male (♂; CeMc) chemical connectome networks, revealed greater asymmetry in males (Fig. 8D; higher *f*^*u*^), and consequently, reduced Redundancy (Fig. 8D; lower *R*^1^). However, the increased asymmetry in the male network was not sufficient to enhance Reachability, which was also lower compared to the hermaphrodite network (Fig. 8D; lower *S*^1^). This was likely due to a difference in relative connectivity between the networks (Fig. 8D; lower male *f*^*d*^), with the male connectome appearing sparser, an effect that can substantially reduce Reachability (Fig. 7E). We observed a similar relationship between chemical and electrical connectome networks both in hermaphrodites and in males (CeH0c and CeMc vs. CeH0e and CeMe). Electrical synapses, which are less abundant than chemical synapses, showed more asymmetry, lower Redundancy, but also lower Reachability (Fig. 8E).

In *C. elegans*, in addition to the main somatic connectome, a small (20 neuron) separate pharyngeal connectome regulates the pharynx, the feeding apparatus of the nematode^32^. We found dramatic differences between the main (CeH0c) and pharyngeal (CePc) chemical networks (Fig. 8F). The pharyngeal network is considerbly denser in its relative number of synaptic connections (Fig. 8F; higher *f*^*d*^), resulting in greater symmetry (Fig. 8F; lower *f*^*u*^) and thus, more Redundancy (Fig. 8F; higher *R*^1^), but also more Reachability (Fig. 8F; higher *S*^1^), likely due to the high volume of connectivity. These feautres may be suited to the particular task of the pharyngeal network, tasked with coordinating food intake.

Finally, we compared network lateralization between different *C. elegans* larval development stages^21^, L1 through L3 (CeL1c, CeL2c, CeL3c). We found a gradual increase in symmetry with development (Fig. 8G; decreasing *f*^*u*^), followed by a return to higher asymmetry in adulthood (Fig. 8G; *f*^*u*^). At the same time, the developing networks exhibited remarkably stable Redundancy (Fig. 8G; *R*^1^), but a steady increase in Reachability (Fig. 8G; *S*^1^), reminiscent perhaps of the progressive interconnectedness appearing in the human brain as it matures^33,34^.

## Discussion

We have shown how the relative level of symmetry vs. asymmetry in synaptic connectivity may settle an inherent tradeoff between Redundancy and Reachability: the selective boosting of specific class paths vs. the expansion and diversification of neuron class connections. We identify a possible tuning principle between symmetry and asymmetry, and thus between Redundancy and Reachability. We find that while contralateral neuron members of each neuron class display symmetry in the number of synaptic contacts they possess (consistent with observed structural symmetry^30,31^), the identity of a subset of these synapses, the *unpaired* synapses, varies between left and right. The number of such unpaired synapses in each neuron is proportional to that neuron’s total number of synapses, and their composition appears to be stochastic. Thus, each neuron may be allocated a certain number of synaptic partners, to which it can randomly link, independent of its contralateral class member, contributing in this way to overall network asymmetry. This allowance of random synaptic connections within each neuron, could enhance Reachability by presenting new possibilities for connectivity.

At the same time, the paired synapses in each neuron show much less stochasticity in their identity, with possible coordination between left and right neuron class members. These synapses are a source of network symmetry, supporting Redundancy by the duplication of selected connections, reinforcing these synaptic paths. Synapse formation and specification involve complex interactions of genetic^23^, biomechanical^24^, activity-dependent^25^ and stochastic^26^ processes. One prediction of our model is that these factors may segregate according to synaptic subpopulation, with paired synapses tending to be prespecified, while unpaired synapses being more stochastic. Evidently, many additional factors may account for neuronal and synaptic lateralization^7^.

Our study focused mostly on the *C. elegans* hermaphrodite connectome, and in particular on chemical synaptic connectivity. However, the principles we have derived seem to hold both in artificially-constructed networks and in other *C. elegans* connectomes, such as in the male, larval and pharyngeal networks. It is notable that although nematodes lack any clear anatomical lateralization in their body, eliminating the need to control left and right extremities or movement, they still exhibit a largely bilateral symmetrical nervous system, with a majority of neurons paired into left-right classes^19^. This suggests that bilateral symmetry may be a deeply conserved feature of the nervous system^7^. Our analysis did not take into account such important features of the connectome as the weight (number or efficacy) or sign (excitatory vs. inhibitory) of each synapse. Some of this information is not yet fully resolved.

A key premise underlying our analyses is that neurons pertaining to the same class share similar or related functions. This is often taken as an implicit assumption in the study of *C. elegans* and other neural circuits. Notably, left/right differences in molecular composition have been identified in several *C. elegans* neuron classes^35^, and functional lateralization, in particular, has been extensively characterized in two specific chemosensory neuron classes, ASE^36–38^ and AWC^39,40^. However, despite the differences between the left and right ASE and AWC neurons, for example, both members of these neuron classes are involved in similar tasks and play related roles in the circuit. In addition, as we have noted, at least half of *C. elegans* neuron classes show chemical and/or electrical synaptic coupling between their bilateral members. Therefore, it is reasonable to assume that analogous information is sent or received by the distinct synaptic contacts of each class member, justifying the neuron classes as a unit for analysis of lateralization.

In more complex systems neuron classes comprise many more than just one bilateral pair of neurons, and the number of synaptic contacts of each neuron can be several orders of magnitude larger than in *C. elegans*. Thus, in such systems an entire brain region could be considered equivalent to a *C. elegans* neuron class. A classic example is the human inferior frontal gyrus (Broca’s area). In the dominant (typically left) hemisphere, this brain region is responsible for language comprehension and speech production^41^, as opposed to the analogous contralateral region that controls prosody. These bilateral functional disparities are associated with left-right dissimilarities in circuit structure and wiring^42,43^. Despite the dramatic differences between these two regions, they are nevertheless bilaterally symmetrical in their overall anatomy and position, they are interconnected by commissural fibers, and ultimately, are both involved in language-related processes. Thus, larger, more complex connectomes may also be amenable to analysis using the Redundancy vs. Reachability framework that we have developed.

In order to properly grasp the impact of lateralization on neuronal structure-function relations, connectome maps must label bilaterally equivalent neurons or structures and resolve the differences and similarities in synaptic connectivity between them, as in the *C. elegans* connectome. In any case, synaptic connectivity should not be assumed to be symmetrical, and such assumptions are unsuitable for validating proper connectome mapping.

As we have shown, it is possible to adapt standard concepts and tools from graph theory to the special case of neural networks, accounting for the grouping of neurons into functional classes with bilateral structure. To this end it is important to distinguish between standard graph paths connecting individual neurons, and realizable class paths that capture an interesting aspect of the interplay between structure and function in neural networks. As noted, our analysis is based on several apparent simplifications, focusing on undirected, unweighted, unsigned exclusively chemical connections. However, the conceptual framework we developed could readily apply to more detailed network representations, enabling further investigation into the impact of synaptic lateralization on salient network properties. Moreover, features like Redundancy and Reachability point at potential functional implications, and at the possibilities afforded by network configuration. In reality, however, neural information flow and dynamics may be much more complex and not necessarily compatible with physical connectivity^44^. When the full complexity of neural networks is considered, effective network function might diverge from that suggested by synaptic connectivity alone. For example, broad long-distance neuropeptide signaling^45,46^ may boost Reachability even in a relatively symmetrical network. Concomitantly, various forms of synaptic plasticity continuously modify the connectivity map. Does plasticity act symmetrically or unilaterally? Can it shift the balance between Redundancy and Reachability or is their impact restricted to a fixed setpoint? A recent study showed a striking switch in lateralization of synaptic connections in the *C. elegans* gustatory circuit following salt learning^47^. Probing the interplay between lateralization and network function prompts many such essential questions.

The broader impact of Redundancy and Reachability, as we have framed these concepts, on overall brain function and, in particular, on such capacities as perception, action, decision making, learning and more, is challenging to discern. Changes in brain lateralization have been shown to be associated with cognitive performance^48^. It can generally be postulated that Redundancy may be especially important for proper sensory or motor function, where robustness and fault tolerance are crucial. Redundancy in synaptic connectivity between specific pairs of neurons may also facilitate learning at the neuronal level^49^. At the same time, Reachability may facilitate the integration, distribution and coordination of neural information, presumably required for more complex brain operation. Interestingly, disease conditions such as schizophrenia have been associated with abnormal connectivity^17^, including a decrease in the level of asymmetry normally observed in healthy brains^50^, and increased segregation in network topology^51^, which could be interpreted as reduced Reachability. Such coupling between reduced asymmetry and Reachability is reminiscent of our findings on the relationship between lateralization and network properties.

## Methods

### Neural networks used in the study

Our study included three types of neural networks, generated and analyzed using the NetworkX library in Python. (1) *C. elegans* connectome networks (Table 1), based on datasets curated to a large part by the Emmons lab (https://wormwiring.org). (2) Artificial neural networks (Table 2), related to the *C. elegans* connectome networks. (3) Symmetrized or desymmetrized networks.

**Table 1.**
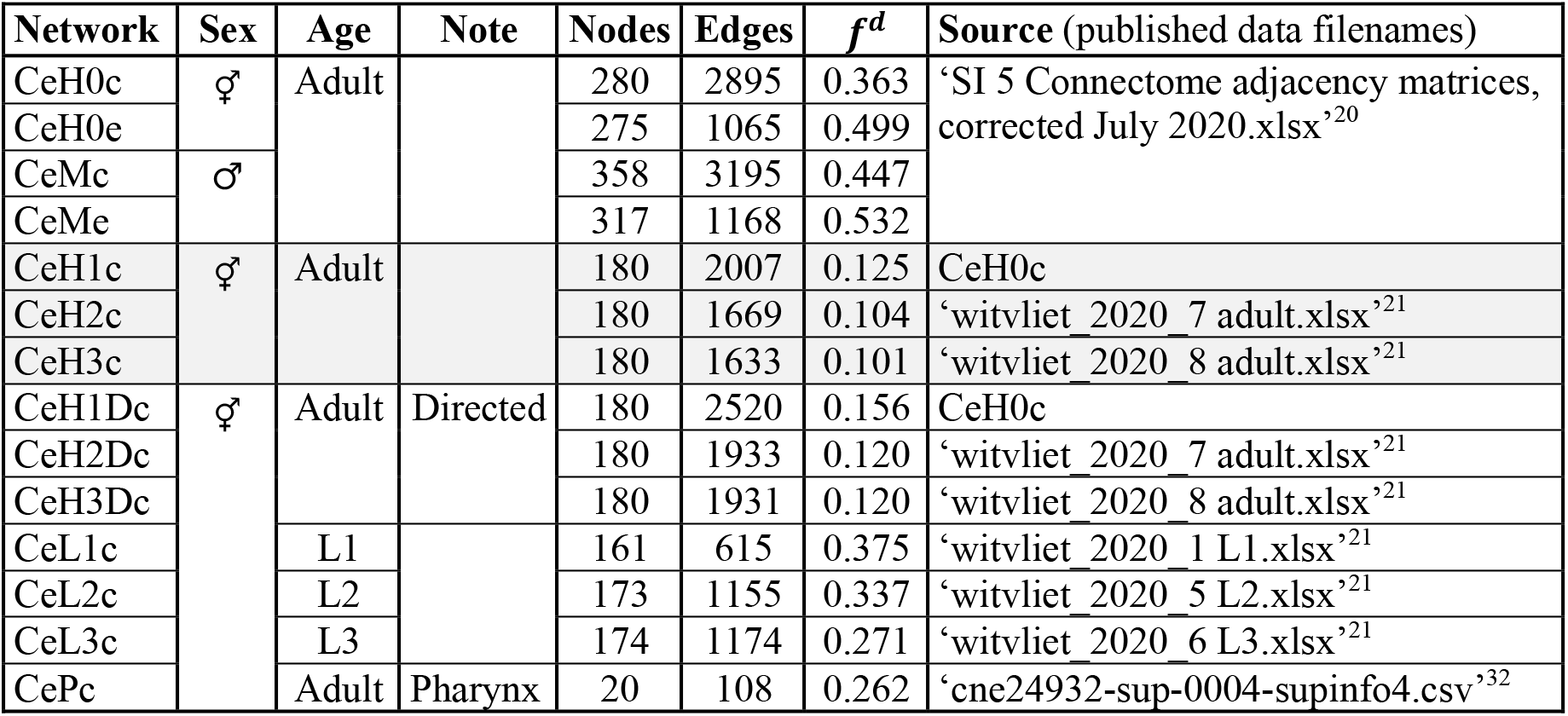
List of *C. elegans* connectome networks. Shaded networks are used throughout most of the study. Unshaded networks used in Fig. 8.

**Table 2.** List of artificial networks and the figures, in which they appear.

| Network | Nodes | Classes | Edges | $f^d$ | Construction method | Figures |
| --- | --- | --- | --- | --- | --- | --- |
| Model | 16 | 8 | 19 | 0.158 | Manual | Fig. 2A, 4A, 5A-C,E,F |
| Small1 | 40 | 20 | 80 | 0.103 | NetworkX: erdos_renyi_graph | Fig. 4B |
| SR | 180 | 68 | 1770 | 0.110 | NetworkX: gnm_random | Fig. 6C,E,F |
| ER | 180 | 68 | 2027 | 0.126 | NetworkX: erdos_renyi_graph | Fig. 6C,F |
| DCB | 180 | 68 | 1994 | 0.124 | Custom |  |
| Small2 | 40 | 20 | 39-156 | 0.05-0.2 | NetworkX: gnm_random | Fig. 7E, S3 |

### Connectome networks

The connectome networks used in this study are listed in Table 1. The shaded networks are the main ones featured in most of the work. The rest of the networks are used in Fig. 8. Unless otherwise specified, all connectome networks were constructed as undirected and unweighted networks, including only chemical synapses. All non-neuronal cells and their synaptic connections were removed. For each network, except for CePc, we also excluded the pharyngeal neurons. Finally, for our analysis, we used the single largest component, the largest connected subgraph within the entire graph, of every connectome dataset, to avoid split networks with disconnected components.

To enable comparison and correlation between different adult *C. elegans* connectome networks, we reduced CeH0c, to a smaller subnetwork, CeH1c, containing the same neurons as CeH2c and CeH3c. Directed CeH1Dc, CeH2Dc and CeH3Dc networks maintained the original directionality assignments of the edges, and allowed for reciprocal edges between the nodes.

### Artificial networks

The artificial networks generated for the study are listed in Table 2. First, the Model network variants, used for demonstration purposes, were composed manually. Second, SR (simple random) networks, as well as the Small2 networks were generated using the ‘gnm_random’ function in NetworkX. This procedure accepts two arguments, *n*, the number of nodes and *m*, the number of edges. For the SR networks, *n* equaled 180, and *m* equaled the average number of edges in the connectome networks (CeH1c, CeH2c and CeH3c), 1,770. The nodes were labeled according to the connectome classes (some of which have more than 2 members). To construct the Small2 networks with variable *f*^*d*^, we varied the number of edges, symmetrized or desymmetrized (see below) the networks and selected networks with specific *f*^*u*^ values.

Third, the ER (Erdős–Rényi) network was generated using the ‘erdos_renyi_graph’ function in NetworkX. The procedure accepts two arguments, *n*, the number of nodes and *p*, the probability of edge creation between any two nodes. *n* equaled 180, and *p* equaled the relative degree, *f*^*d*^ = 0.125, of the undirected *C. elegans* connectome network CeH1c. To generate the set of undirected vs. directed Small1 networks, we first constructed 20 networks using the ‘erdos_renyi_graph’ function with *n* = 40, *p* = 0.1. We applied symmetrization and desymmetrization (see below) to derive 3 sets of 20 networks each with a different average *f*^*u*^ value for each set. We then created for each undirected network an equivalent directed network by randomly assigning directionality to every edge. This was done using the following algorithm:

#### Algorithm 1: Convert undirected to directed

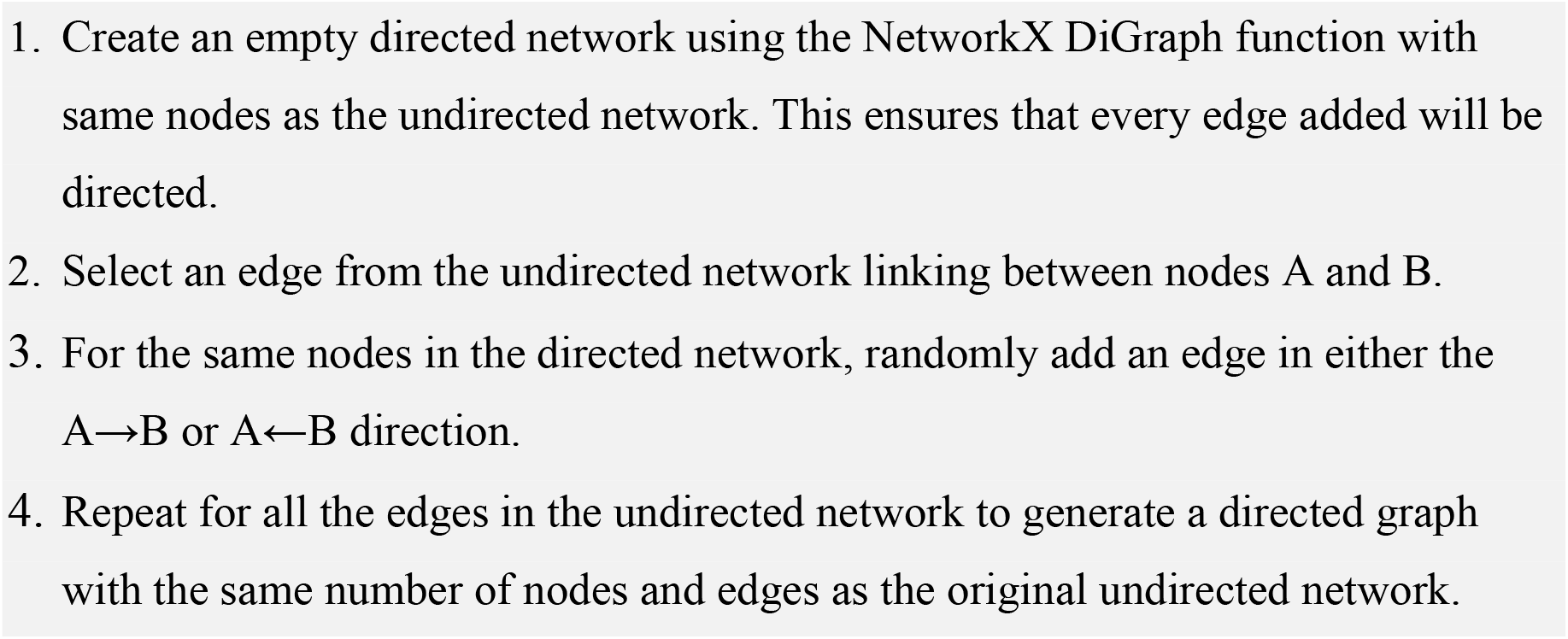

Finally, the DCB (degree connectome-based) network was generated as a random network with a degree distribution similar to that of the connectome network CeH1c. First, we created a network with 180 nodes and no edges. Then for each node, edges were formed with random partners out of a list combining the partners of that node in the CeH1c network and in the ‘contactome’, the physical adjacency between each two neurons in the connectome^30,31^ (we used the source file ‘cel_n2u_nr_adj.csv’^31^). This process was repeated until each node reached its target degree.

As in the connectome networks, when generating the artificial networks, we only accepted network outputs where all the nodes belonged to the single largest component (the largest connected subgraph within the entire graph).

### Symmetrized and desymmetrized networks

In order to construct networks ranging from fully symmetrical to fully asymmetrical, we developed an algorithm for symmetrizing or desymmetrizing an initial network (Fig. S4). The algorithm is based on iterative addition and removal of edges in the network to progressively reduce or increases *f*^*u*^, while maintaining network degree. We applied this algorithm to initial networks from Table 1,2, and obtained in this manner sequences of networks with incremental symmetry/asymmetry levels.

#### Algorithm 2: Network symmetrization / desymmetrization

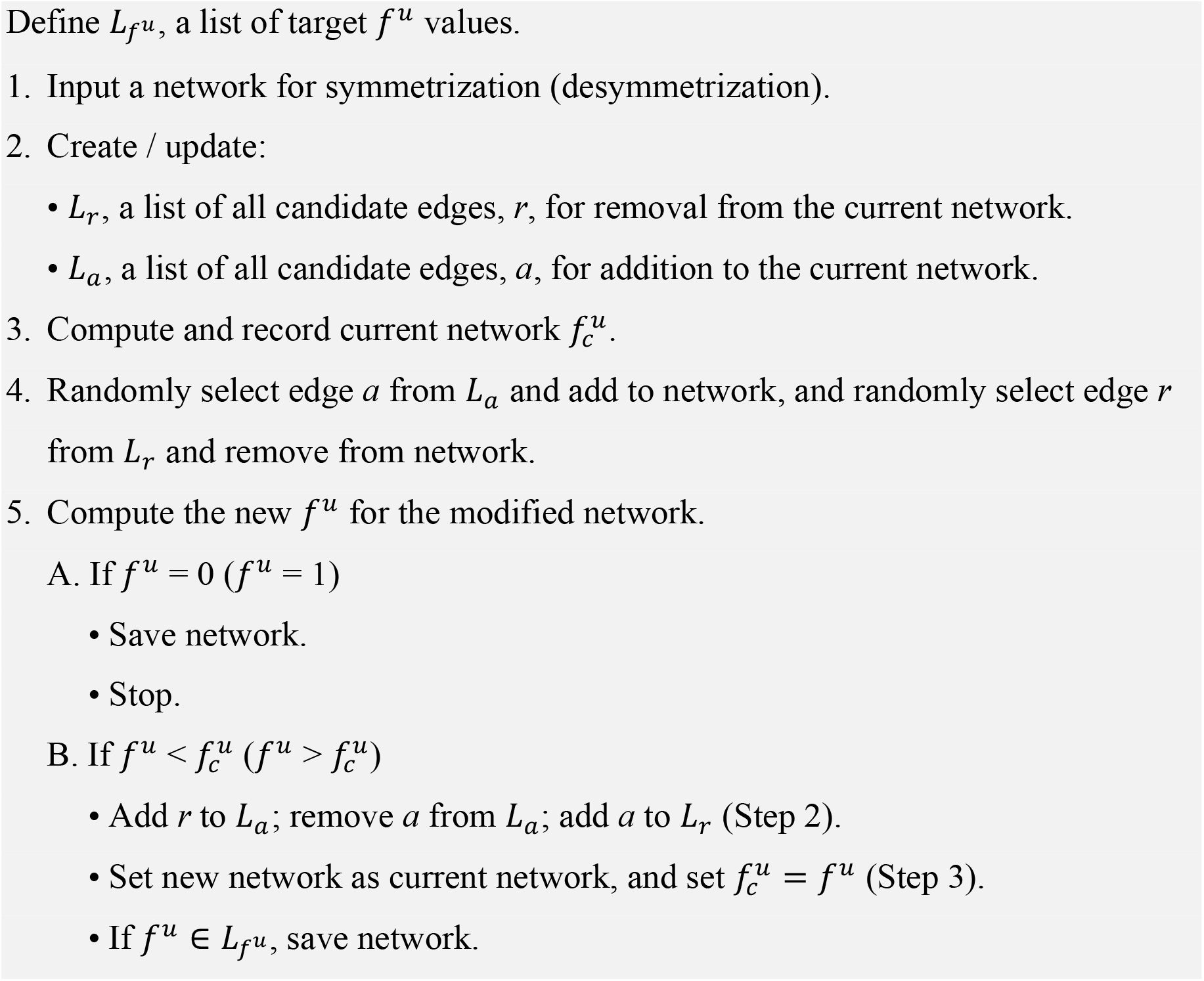
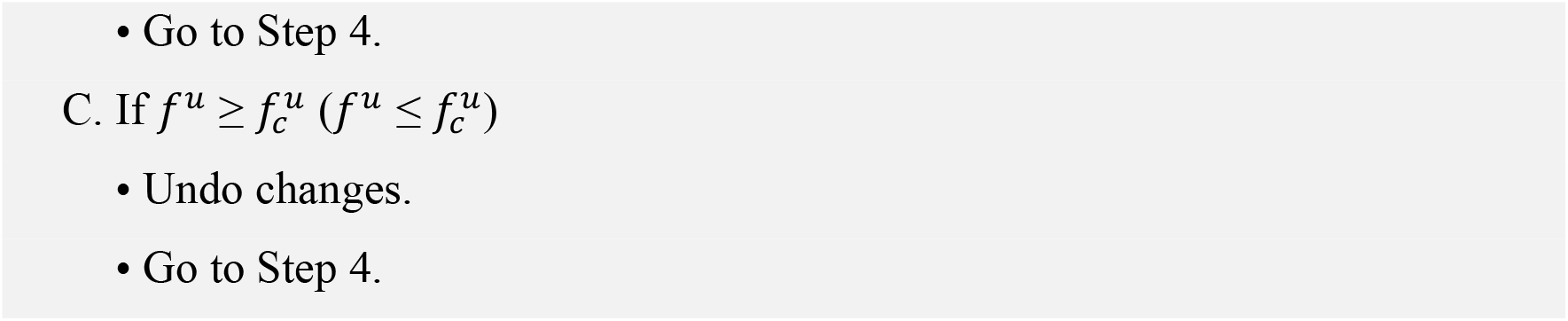

### Computing symmetry/asymmetry metrics

We developed custom algorithms for computing symmetry and asymmetry metrics. All functions are from NetworkX. First, in order to compute the Redundancy metric, *R*^*n*^, it is necessary to derive 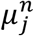, the number of neuron paths of length *n* that realize each class path, *j*. The computation considers all neuron classes, including those with more than 2 bilateral neuron pair members. ^★^The 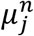 algorithm is also used for deriving Redundancy based exclusively on paired synapses, *R*^*p*,1^ (in the special case of *n* = 1). In this case, only neuron classes comprising a bilateral pair of neurons are included. Thus, *R*^*p*,1^ represents a lower bound for the contribution of paired synapses to Redundancy.

#### Algorithm 3: *R*^*n*^ Redundancy (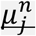 and *R*^*p*,1^)

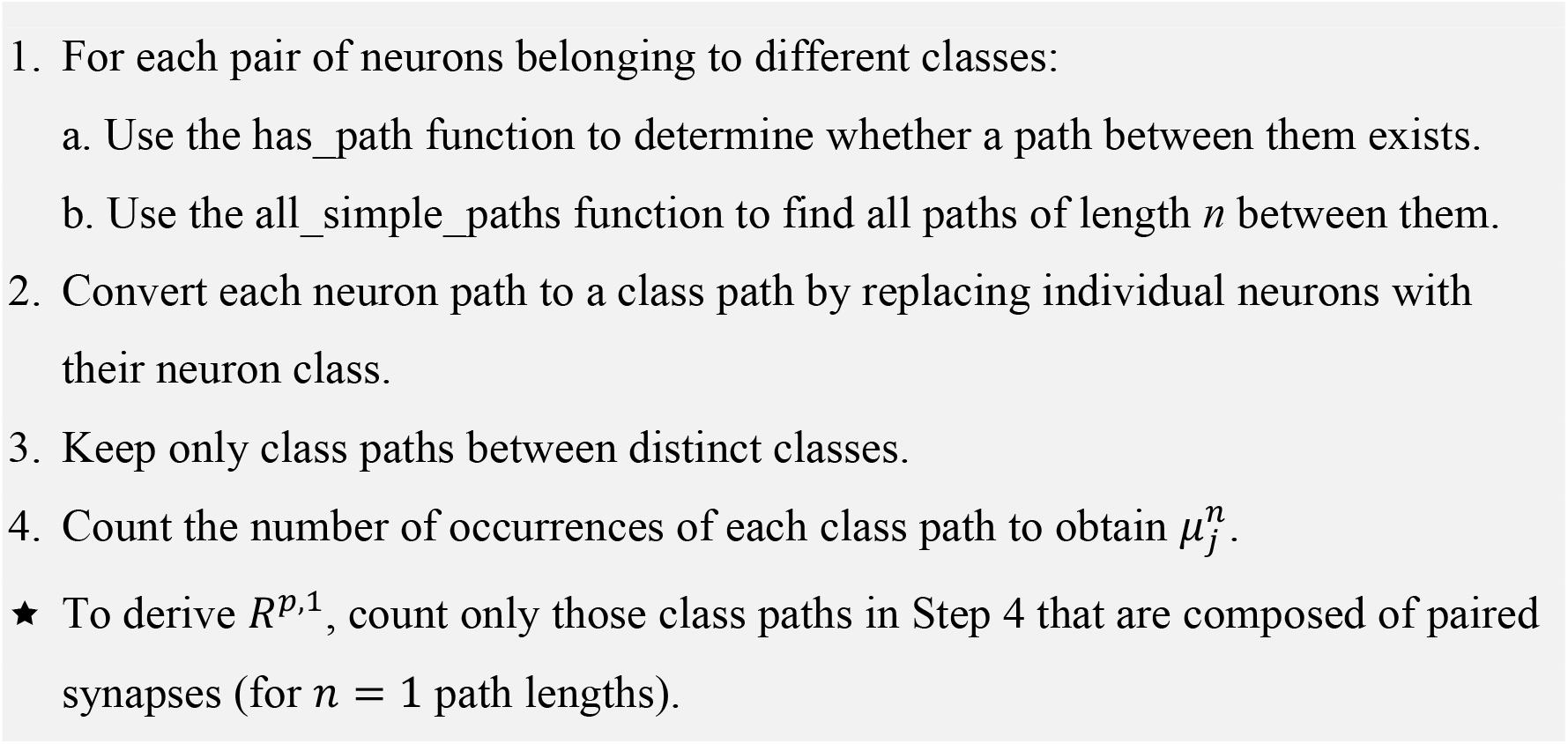

Second, to compute the Reachability metric, *S*^*n*^, it is necessary to first derive 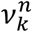, the number of neuron classes that can be reached from each neuron class, *k*, by a realizable class path of length, *n*. This computation includes also neuron classes with more than 2 bilateral neuron pair members. ^★^The 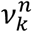algorithm is also used for deriving Reachability based exclusively on unpaired synapses, *S*^*u*,1^ (in the special case of *n* = 1). In this case, only classes composed of 2 bilateral neuron pairs are considered.

#### Algorithm 4: *S*^*n*^ Reachability (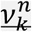 and *S*^*u*,1^)

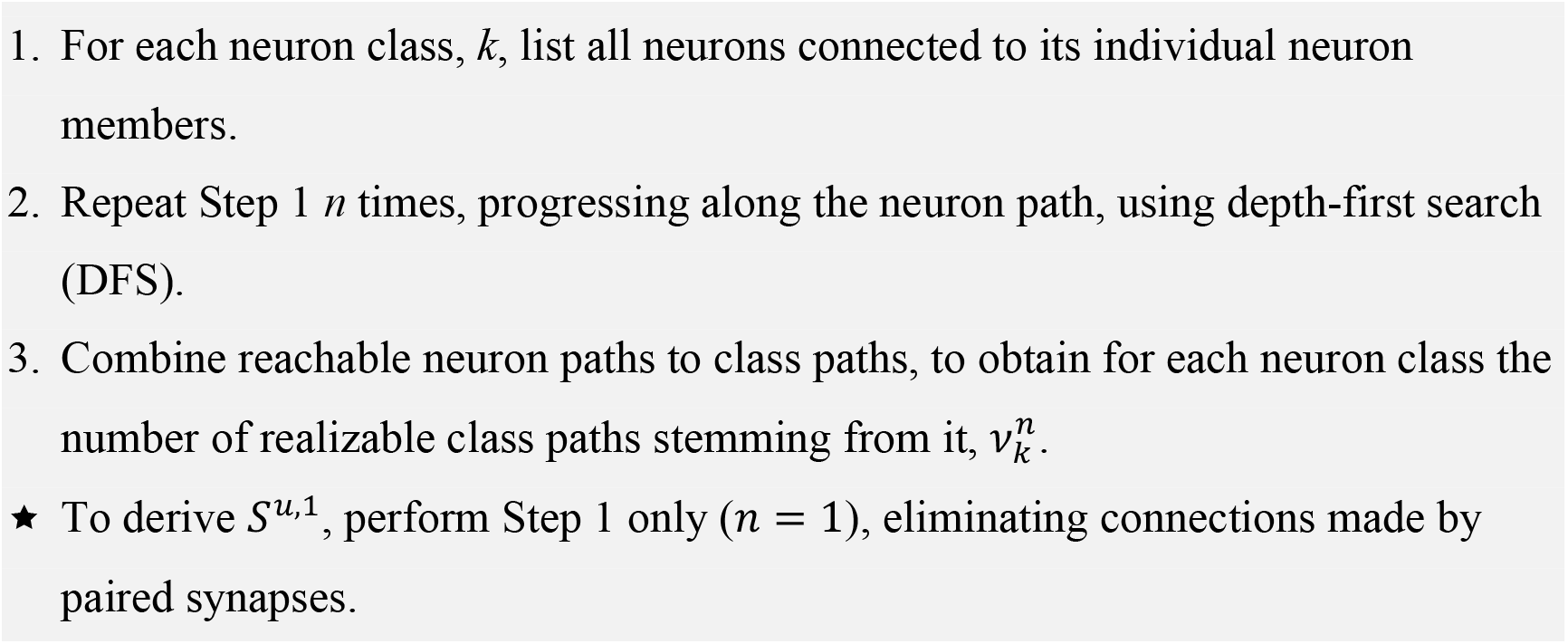

## Acknowledgments

The study was funded by Israel Science Foundation Grant 1465/20 and by the Horizon Europe, PathFinder European Innovation Council Work Programme under grant agreement No 101098722. Views and opinions expressed in this article are those of the authors only and do not necessarily reflect those of the European Union or European Innovation Council and SMEs Executive Agency (EISMEA). Neither the European Union nor the granting authority can be held responsible for them.

## Author contributions

Conceptualization, V.S.B and I.R.; Methodology, V.S.B and I.R.; Software, V.S.B.; Formal Analysis, V.S.B and I.R.; Investigation, V.S.B and I.R.; Writing – Original Draft, I.R.; Writing – Review & Editing, V.S.B and I.R.; Visualization, I.R.; Supervision, I.R.; Funding Acquisition, I.R.

## Supplemental information

Document S1. Figures S1-S4.

Document S2. Source data.

