## Supplemental Figures for "Asymmetry in synaptic connectivity balances redundancy and reachability in the *C. elegans* connectome"

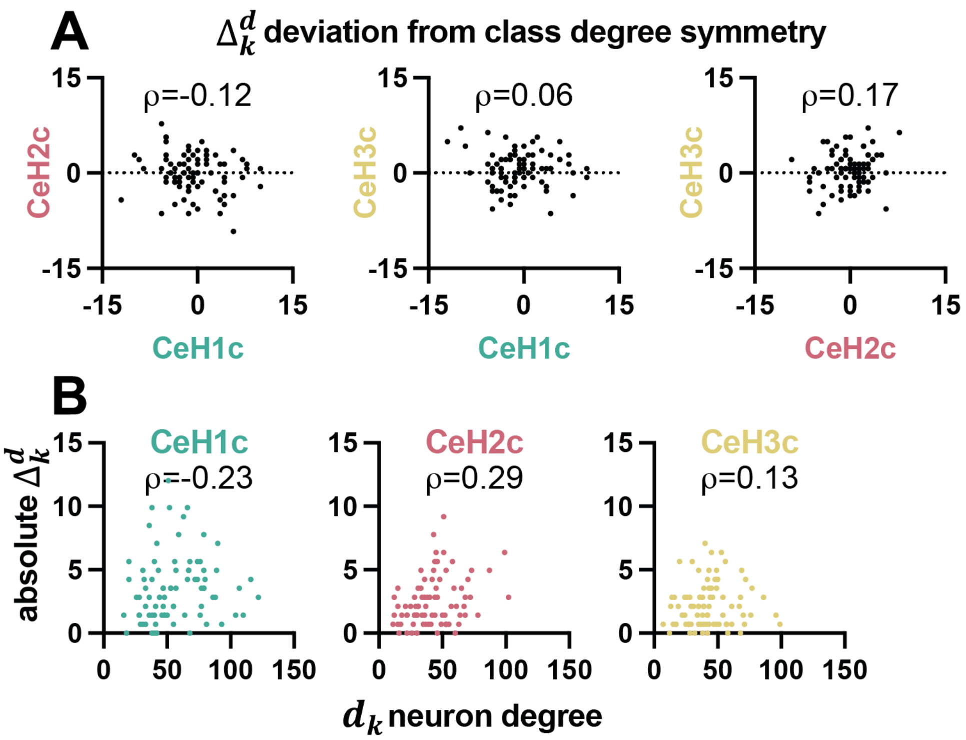


**Figure S1** Deviations from left-right symmetry in neuron chemical synapse degree.

(A) Comparison of $\Delta_k^d$ between connectome datasets. $\rho$ - Spearman rank correlation. n=83 for all plots. p=0.2823, 0.5597, 0.1273, respectively.

(B) Comparison between the absolute value of $\Delta_k^d$ and neuron class degree, $d_k$, for each connectome dataset. $\rho$ - Spearman rank correlation. n=83 for all plots. p=0.0384, 0.0082, 0.2530, respectively.

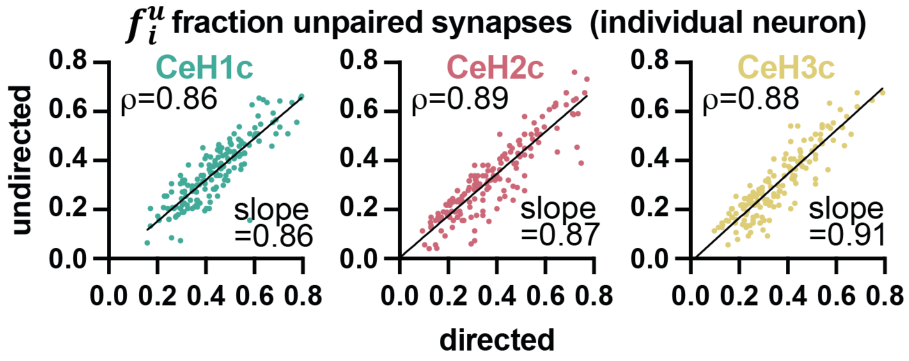


**Figure S2** Comparison of the fraction of individual neuron, $f_i^u$ unpaired synapses, between undirected and directed connectome networks. $\rho$ - Spearman rank correlation coefficient. Slope as fitted by a linear regression. n=166, p<0.0001 for all plots.

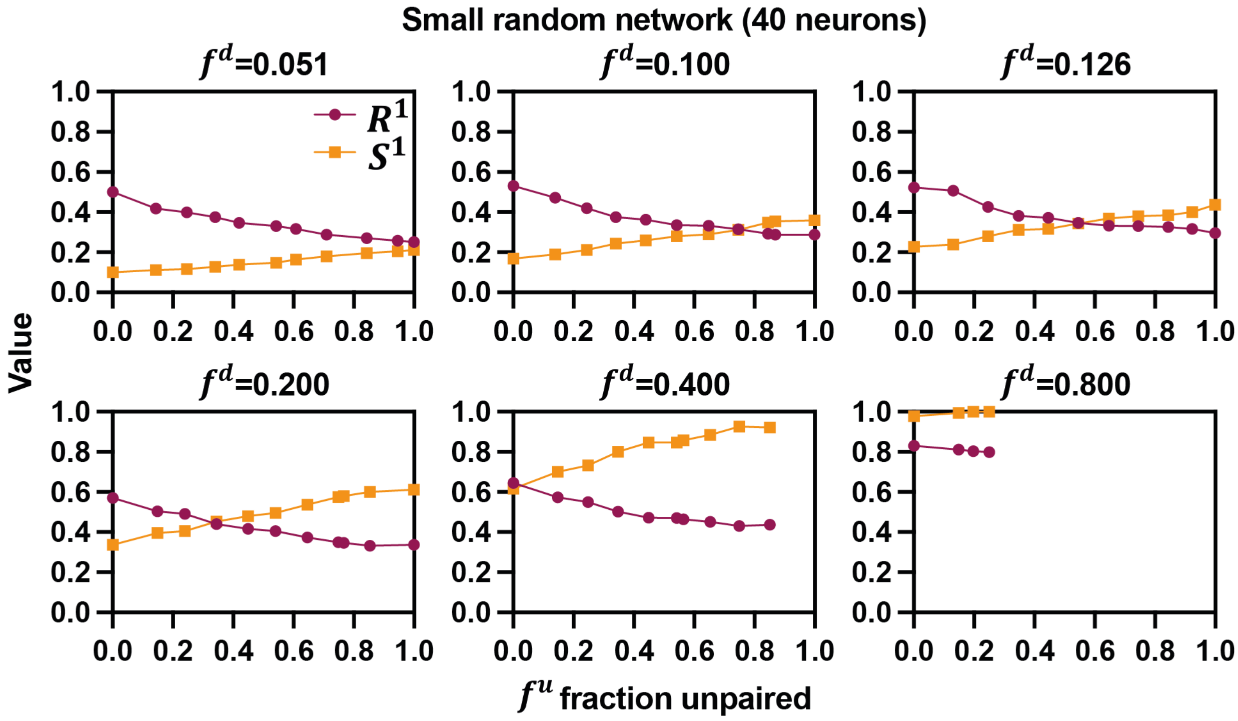


**Figure S3** Relative network degree affects the relationship between $R^1$, $S^1$ and $f^u$. Redundancy, $R^1$, and Reachability, $S^1$, of a series of small random networks (40 neurons) with varying number of synapses ($f^d$) plotted against the fraction of unpaired synapses, $f^u$ of each network.

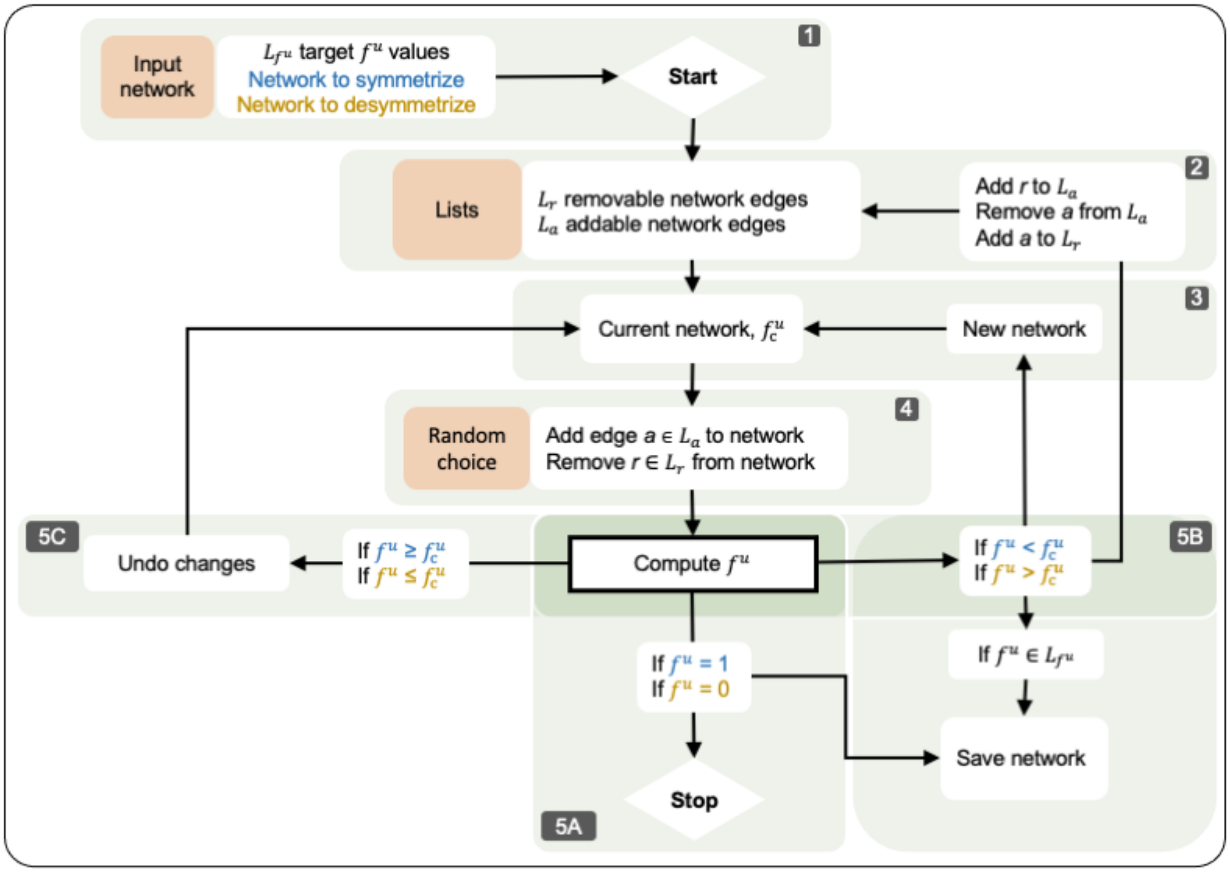


**Figure S4** Schematic flow chart of symmetrization / desymmetrization algorithm (see Methods).
